# Treating the untreatable: Reversing β-lactam resistance in MRSA by membrane-domain-dissolving antibiotics

**DOI:** 10.1101/2025.10.27.684379

**Authors:** Adéla Melcrová, Willem Woudstra, Michaela Wenzel, Mariella Gabler, Wenche Stensen, John S. M. Svendsen, Wouter H. Roos, Romana Schirhagl

## Abstract

Antimicrobial resistance is one of the most serious threats to global health with methicillin-resistant *Staphylococcus aureus* (MRSA) being the leading Gram-positive pathogen. Recently we discovered a new antimicrobial target for small antibiotics — dissolution of functional domains in the bacterial membrane without pore formation. Here we apply this mechanism to reverse β-lactam resistance in MRSA. We show that supplementation with a membrane-domain-dissolving antibiotic lowers the susceptibility of both a MRSA clinical isolate and the methicillin-susceptible *S. aureus* (MSSA) to multiple β-lactams several folds. In particular, full reversal of resistance to oxacillin and penicillin in MRSA is achieved. We provide evidence on the nano-scale (atomic force microscopy), living bacteria (fluorescence microscopy), and bacterial cultures (microbiology assays) that the β-lactam resistance reversal is linked to dissolution of membrane domains. A general nature of this principle is expected, and could be applied using various membrane-domain-dissolving antibiotics to reverse β-lactam resistance in various bacteria.

TOC Figure:
Model of β-lactam susceptibility renewal in MRSA based on data presented in this study.

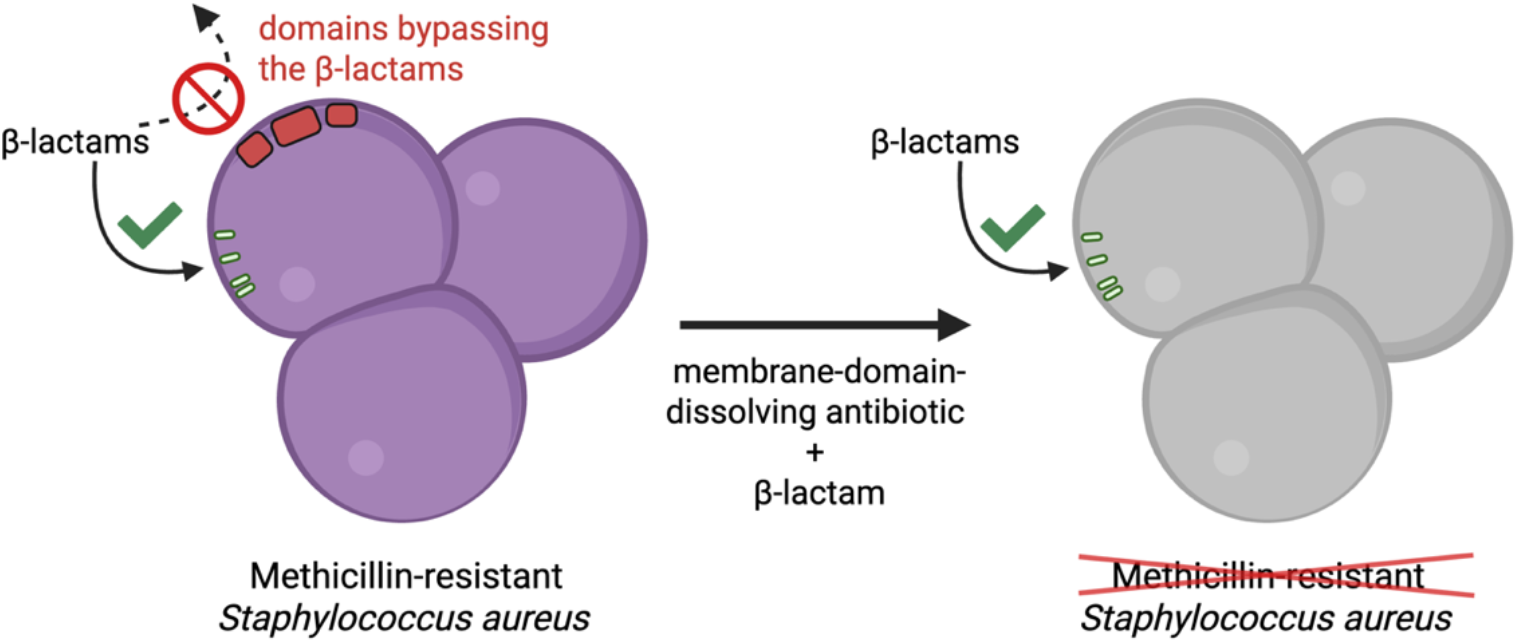

## INTRODUCTION

Antimicrobial resistance has been declared as one of the top ten threats to global health by the World Health Organization (WHO)^1^. According to a comprehensive WHO study on the spread of antibiotic-resistant bacteria published in 2024^2^, methicillin-resistant *Staphylococcus aureus* (MRSA) and other resistant *cocci* are high-priority pathogens in urgent need of effective treatment options. The present work describes a new principle that reverses β-lactam resistance in MRSA, thus reenabling the effective use of classical antibiotics like oxacillin.

Recently, we described a novel antibiotic mechanism of action, the dissolution of nanoscopic domains in the bacterial membrane^3^. During this process, antibiotic molecules insert themselves into the outer membrane leaflet of the bacterial cytoplasmic membrane, dissolving the natural domain distribution of the native lipid mixture. Such membrane-domain-dissolving activity, resulting in membrane homogenization, was first observed for the peptide-like molecule AMC-109^3^ and later also confirmed for lipopeptide N-alkylamide 3d^4^. In both cases, we were able to directly visualize this phenomenon on the nanoscale and at sub-second imaging speed using high-speed atomic force microscopy (AFM) of lipid membranes extracted from methicillin-sensitive *S. aureus* (MSSA). Although chemically different, both agents feature a size comparable to a single phospholipid molecule and are partially cationic and partially hydrophobic. Hence, they prefer to locate at the interface between the hydrophobic interior and the negatively charged headgroup region of the bacterial cell membrane. Importantly, these membrane-domain-dissolving antibiotics effectively homogenize the membrane contents without causing further damage in the form of pores or perforations, which makes these molecules distinct from most other antimicrobial peptides.

The mechanism of action of β-lactam antibiotics is based on blocking the synthesis of the peptidoglycan cell wall that surrounds the bacterial cell membrane. This is achieved by binding to and inhibiting the enzymatic activity of penicillin-binding proteins (PBPs), which cross-link the freshly synthesized peptidoglycan subunits into a stable mesh^5–8^. The resistance of *cocci* to both penicillinase-sensitive and non-sensitive β-lactams is often associated with the presence of the low-affinity PBP2a^8–11^. This specific PBP has its active site located in a deep pocket that cannot be reached and blocked by β-lactams^11,12^. Consequently, PBP2a-expressing bacteria are able to continue peptidoglycan cross-linking in the presence of β-lactam antibiotics. However, even in MRSA, β-lactams still block all other available PBPs making the peptidoglycan synthesis possible but not ideal^11^.

Several studies^13–16^ have shown that PBP2a is located in functional membrane microdomains (FMMs) in the bacterial cell membrane and that the disassembly of these microdomains reverses MRSA resistance to β-lactams. Hui et al.^13^ achieved FMM disassembly with pulsed laser treatment. Yet, this treatment also damages the surrounding membrane and makes it porous. García-Fernández et al.^16^ proved that MRSA mutants that do not assemble FMMs at all are no longer β-lactam-resistant. So far, no disassembly of FMMs in an unmodified, living MRSA has been achieved without inducing further membrane damage. AMC-109 and N-alkylamide 3d ^3,4^ are capable of specifically disassembling FMMs, potentially opening up new avenues for resistance-reversing combination therapy approaches.

Their membrane-domain-dissolving mechanism has previously been demonstrated on lipid membranes extracted from MSSA^3,4^. However, FMMs in living bacteria are complex, containing not only lipids but also proteins and other molecules. In our previous studies^3,4^, we hypothesized that membrane-domain-dissolving antibiotics do have the capability of dissolving native FMMs in living bacteria. Based on this, we hypothesized further that antibiotic-mediated FMM dissolution could constitute a tool to reverse the resistance of MRSA to *β*-lactams.

In this study, experimental evidence for this hypothesis is provided for the membrane-domain-dissolving agent AMC-109 in combination with three *β*-lactam antibiotics, penicillin G, oxacillin, and methicillin, both in MSSA and MRSA. Penicillin G (or benzylpenicillin) is a natural *β*-lactam antibiotic that was among the first agents used against a variety of Gram-positive pathogens^17^. Some *cocci*, however, produce the enzyme penicillinase (or *β*-lactamase) that inactivates penicillin G^17,18^, conferring resistance. In the 1960’s new synthetic penicillin variants resistant to hydrolysis by penicillinase were synthesized^8,10,19–21^, among them methicillin and oxacillin. Nowadays, methicillin is no longer used for its degradation during storage associated with the high risk of acute kidney damage in patients^17,22,23^. Oxacillin is administered in the form of oral pills and as intramuscular or intravenous injections. For critically ill patients, continuous oxacillin infusions can be used to achieve a resting oxacillin concentration in blood serum higher than 4× minimum inhibitory concentration (MIC) as determined by the international standards^24–26^, i.e. >8 µg/ml.

Here, we show that the partial dissolution of membrane domains by subinhibitory concentrations of a membrane-domain-dissolving antibiotic considerably increases the activity of all three tested β-lactams against both MSSA and MRSA. Importantly, oxacillin and penicillin G MICs were lowered far below their clinical breakpoints, re-sensitizing MRSA to β-lactams and achieving highly effective eradication of MRSA cultures. Using high-speed AFM, fluorescence live cell imaging, and spectroscopic membrane fluidity measurements, we provide evidence for a causal connection between the dissolution of FMMs and reversal of β-lactam resistance. We thus present a novel mechanism underlying the synergy of membrane-domain-dissolving antibiotics with β-lactams with fundamental implications for the design and development of combination therapy approaches against MRSA and other pathogens.

## RESULTS AND DISCUSSION

### Domain-dissolving antibiotic AMC-109 reverses *β*-lactam resistance

Minimum inhibitory and bactericidal concentrations (MIC and MBC) of the *β*-lactams penicillin G, oxacillin, and methicillin, and of the membrane-domain-dissolving antibiotic AMC-109 were determined against both MRSA and MSSA (Table 1). The clinical breakpoint, meaning the concentration cutoff for classifying a bacterial strain antibiotic-susceptible, is ≤0.125 µg/ml for penicillin against *S. aureus*, according to both the European Committee on Antimicrobial Susceptibility Testing (EUCAST)^25^ and the United States Food and Drug Administration (FDA)^26^. Due to the storage instability of methicillin, methicillin resistance is tested using oxacillin^27,28^. The clinical breakpoint for oxacillin against *S. aureus* used to be for a long time considered as ≤0.5 µg/ml, which has been updated in 2021 to ≤2 µg/ml^26,29^. No limits are given for methicillin and AMC-109 as these antibiotics are not currently approved for clinical use.

**Table 1:** MICs and MBCs of MRSA and MSSA after 24 h exposure to penicillin G, oxacillin, methicillin, and AMC-109. Maximum MIC and MBC obtained from 5–14 times repeated experiments are shown.

| Bacteria | MRSA |  | MSSA (ATCC 29213) |  |
| --- | --- | --- | --- | --- |
| | MIC [$\mu\text{g/ml}$] | MBC [$\mu\text{g/ml}$] | MIC [$\mu\text{g/ml}$] | MBC [$\mu\text{g/ml}$] |
| Penicillin G | 2 | 2 | 0.25 | 0.5 |
| Oxacillin | 2 | 2 | 0.5 | 0.5 |
| Methicillin | 8 | 8 | 2 | 2 |
| AMC-109 | 2 | 2 | 4 | 4 |

Table 1 shows that our clinical MRSA isolate is penicillin-resistant as evidenced by the penicillin G MIC >0.125 µg/ml. Our MRSA isolate with its oxacillin MIC = 2 µg/ml is considered borderline methicillin-susceptible according to the new susceptibility standards and resistant according to the clinical breakpoint valid up to 2021. The MSSA strain is sensitive to oxacillin, but is borderline resistant to penicillin G. This result was expected as MSSA ATCC 29213 expresses a β-lactamase^30^ that breaks down the natural penicillin G, but is unable to hydrolyze newer synthetic β-lactams^20^.

Overall, the MSSA strain is 4–8 times more sensitive to all tested *β*-lactams compared to MRSA. The MIC of the membrane-domain-dissolving antibiotic AMC-109 is 2 µg/ml for MRSA and 4 µg/ml for MSSA. With reported toxic concentrations for human cells in the order of ∼100 µg/ml^31^, this leaves a 25 to 50-fold therapeutic window. AMC-109 clearly shows consistently good activity regardless of the β-lactam resistance status, a result that corresponds well to previous reports on AMC-109^32^, showing an MIC range of 2–4 µg/ml and an MBC range of 2–8 µg/ml across 155 clinical MRSA isolates. The MBC values for all tested antibiotics (with one exception of penicillin G against MSSA) are the same as the MIC. This demonstrates that these antibiotics eradicate the bacteria and not only inhibit their growth.

After determining the individual MIC and MBC values, checkerboard assays were used to evaluate the synergistic effects between AMC-109 and the three β-lactams (Figure 1). Figure 1 shows the results of one exemplary experiment for each bacterial strain and AMC-109/β-lactam combination. AMC-109 shows a remarkable potency to lower the dose of β-lactams needed to stop bacterial growth. This can be seen from the substantial shift of growth inhibition to lower β-lactam concentrations at supplementation with 0.5–1 µg/ml AMC-109, which was observed for all tested antibiotic combinations against both MRSA and MSSA. In case of MRSA, the inhibitory concentrations of penicillin G and oxacillin fell deep below the resistance breakpoints when supplemented with 1 µg/ml AMC-109, effectively reversing MRSA resistance. A similar trend was already observed at 0.5 µg/ml AMC-109, albeit with less potency. Across replicate experiments, an AMC-109 concentration of 0.5–1 µg/ml was consistently able to lower β-lactam MICs, while the extent of β-lactam potentiation varied between 2 and 256-fold (Figure S1). MBCs were consistently within one dilution step of their respective MIC values, confirming that all combinations are bactericidal.

**Figure 1:**
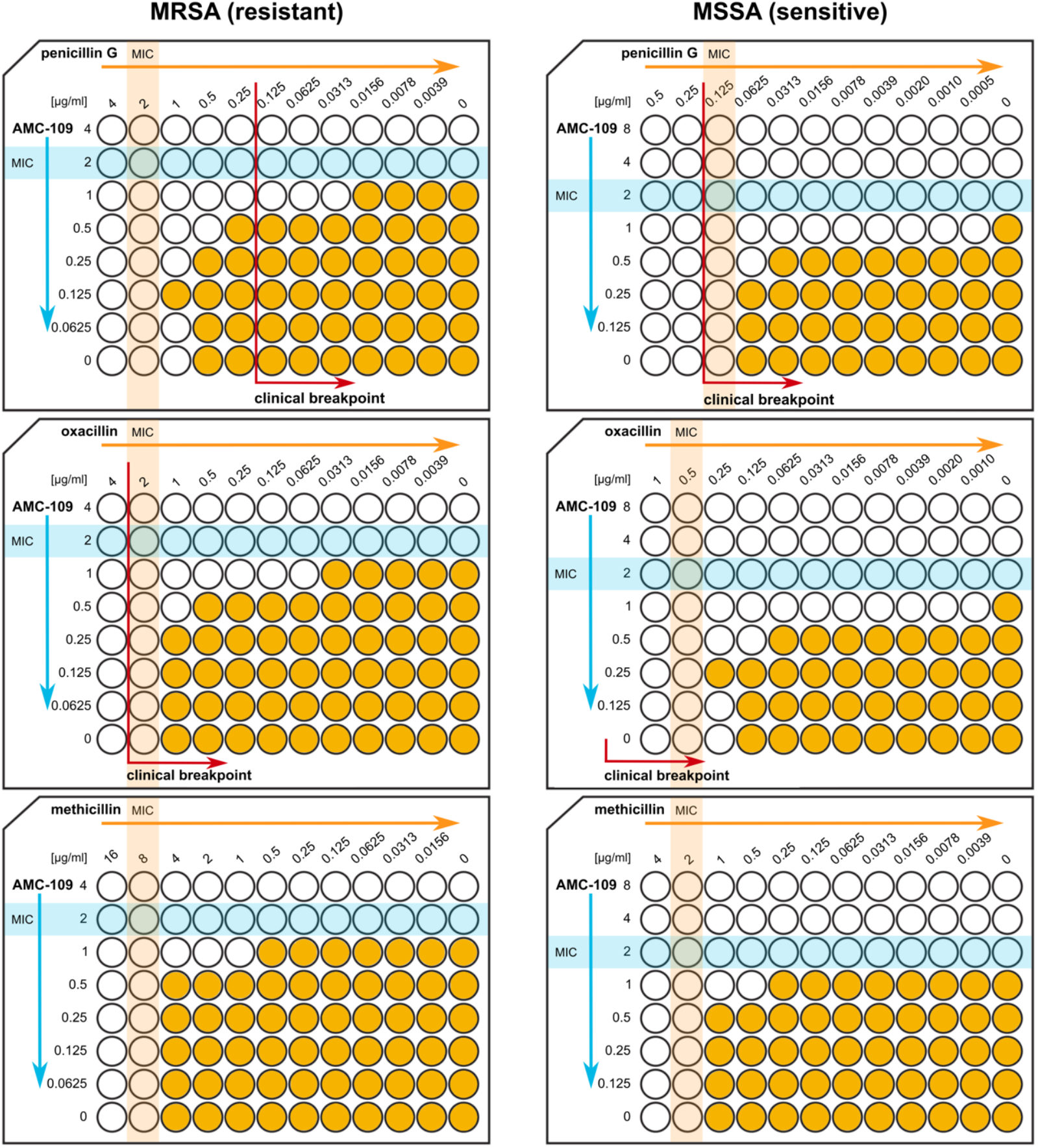
Supplementation with sub-inhibitory concentrations of AMC-109 (0.5–1 µg/ml) increases the sensitivity of MRSA and MSSA to all tested *β*-lactams (penicillin G, oxacillin, methicillin). MIC values for individual antibiotics are highlighted in light blue (AMC-109) and light orange (*β*-lactam). Red arrows denote clinical breakpoints for each *β*-lactam according to FDA and EUCAST regulations^25,26^. Wells with bacterial growth are depicted in dark orange, wells without growth in white. The figure depicts a representative example out of 3–4 replicates performed for each strain and antibiotic combination.

It is noteworthy that the tested combinations resulted in the unidirectional potentiation of β-lactams by AMC-109, and not reciprocal synergy, which would involve the simultaneous lowering of the growth-inhibitory doses of both antibiotics. This observation supports our hypothesis that the dissolution of membrane domains by AMC-109 disrupts the membrane organization required for oligomerization of PBP2a and consequently β-lactam resistance^16^. Bacteria will then be forced to rely on β-lactam-sensitive PBPs, resulting in growth inhibition and cell death.

The fact that a similar potentiation was also observed with MSSA points towards other PBPs being located in or at least impacted by membrane domains^5^ that are dissolved by AMC-109. However, it is also possible that domain dissolution impairs production and/or secretion of extracellular penicillinases, activity of membrane-bound penicillinases, or the signaling cascade that recruits penicillinases to the membrane^33,34^.

### AMC-109/oxacillin combination kills MRSA

For a deeper understanding of the kinetics of the selected antibiotic combination, time-kill assays were performed. Based on the efficacy of the combined treatments in the checkerboard assays, we chose to study in detail the antibiotic effects of the combination of 1 µg/ml AMC-109 together with 0.25 µg/ml oxacillin on MRSA. This oxacillin concentration is below the MIC breakpoint for resistance by the FDA^26^, and appears to have renewed activity on MRSA when supported by 1 µg/ml AMC-109 (Figure 1). Figure 2 shows the average number of colony forming units per ml (CFU/ml) from the 3-times repeated time-kill assays and the associated standard errors of the mean. Untreated cultures and cultures exposed to each antibiotic separately were included as controls. The untreated control (Figure 2, black) grows already after 4 h from ∼10^7^ CFU/ml until a saturated ∼10^8^ CFU/ml. The oxacillin-treated culture (Figure 2, magenta) showed a reduced growth rate compared to the untreated control but no bactericidal effects. In contrast, 1 µg/ml AMC-109 (Figure 2, blue) acted rapidly bactericidal decreasing the CFU/ml count from ∼10^7^ to ∼10^4^ in 4 h, yet allowed for partial recovery of growth after 6h (statistical significance of the growth recovery is p<0.05 at 24h, and p<0.01 at 48h). Interestingly, the combined treatment (Figure 2, red) resulted in bactericidal activity without CFU recovery resulting in CFU/ml reduction from ∼10^7^ to ∼10^3^. This prevention of recovery in the bacterial culture when oxacillin is present, suggests that AMC-109 indeed allows for the oxacillin activity to be restored, which in turn supports the potentiated bactericidal effect of the two antibiotics.

**Figure 2:**
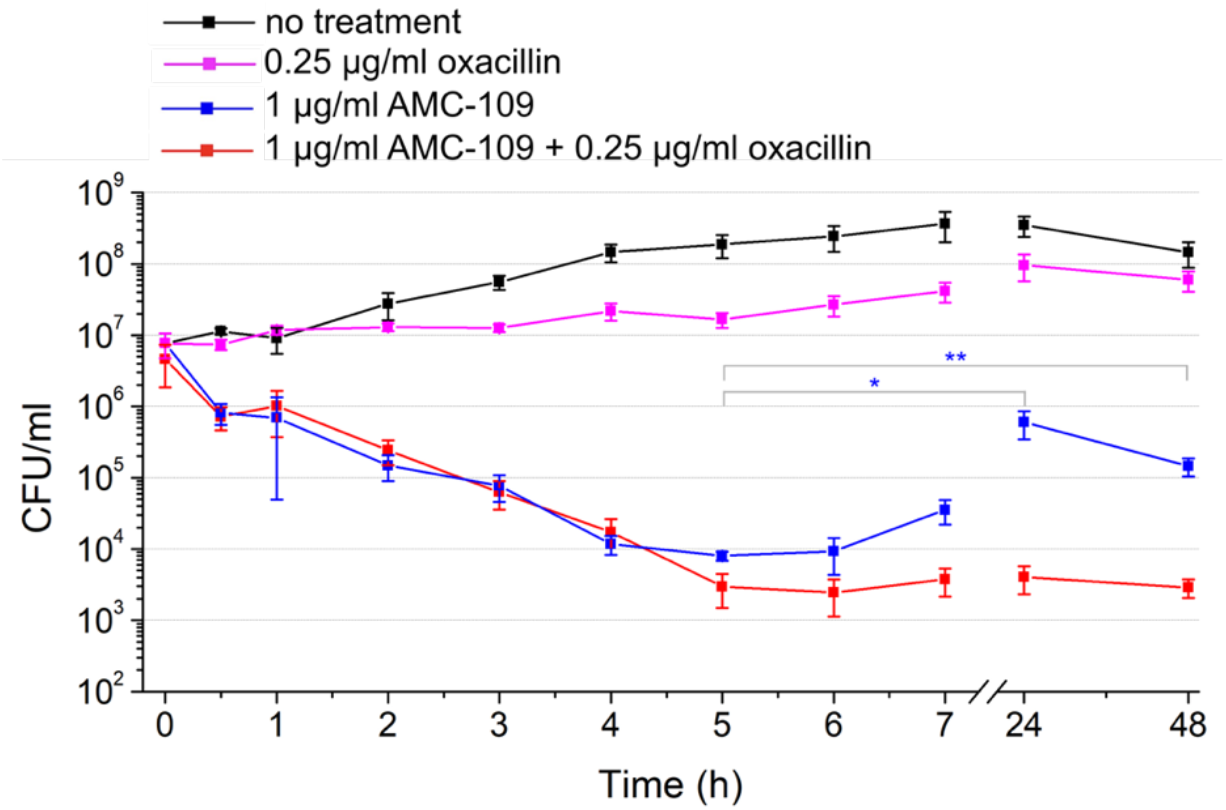
Time-kill assays of MRSA upon exposure to the combination therapy of 1 µg/ml AMC-109 + 0.25 µg/ml oxacillin (red). Control measurements of bacteria exposed to no treatment (black), 0.25 µg/ml oxacillin (magenta), and 1 µg/ml AMC-109 (blue) are provided. Data points are averages of 3 biological replicates. Error bars represent standard errors of the mean of 3 biological replicate experiments (see Figure S2 for individual replicates and Figure S3 for corresponding optical density measurements). Note that although 1 µg/ml AMC-109 initially reduces the bacterial cell count, it is still a sub-inhibitory treatment against MRSA as the colony growth is resumed after the 6 hours. Asterisks denote the statistical significance of the recovered growth of the culture exposed to 1 µg/ml AMC-109; * indicates p-value p<0.05, ** indicate p<0.01. Moreover, the MIC experiments done with a starting concentration of 10^5^ CFU/ml show visible growth after 24 hours (Figure 1).

### Reversal of β-lactam resistance coincides with nanoscopic domain dissolution in MRSA lipid membranes

Using lipid membranes extracted from MSSA, we have previously shown that AMC-109 clusters and consequently dissolves nanoscopic membrane domains at above MIC concentrations^3^. In the combined AMC-109/oxacillin treatment, sub-inhibitory concentrations of the membrane-domain-dissolving antibiotic are used. To test the effects of sub-inhibitory AMC-109 levels on MRSA and MSSA membranes, we prepared total lipid extracts from the same MRSA clinical strain as used in other assays in this study and compared with new experiments on the lipid extracts from MSSA.

High-speed AFM experiments were performed on extracted MRSA lipid membranes supported on a solid substrate. The MRSA membranes were first imaged in phosphate buffered saline (PBS) with an imaging speed of 300–500 ms per frame to observe domain movement. The MRSA lipid membranes contained clearly visible phase-separated domains, which were distinguishable as thicker parts of the membrane (Figure 3A, bright golden spots), and exhibited clear domain movement (Figure S4). The lateral movement of the domains was too fast to track individual domains, even at sub-second imaging speed. Upon addition of 0.5 µg/ml AMC-109 (Figure 3B), the domains slow down their movement after a few minutes and cluster together. No domain dissolution was observed at this concentration. After increasing the AMC-109 concentration to 1 µg/ml, the clustering effect was more pronounced and partial domain dissolution was observed. Figure 3C show the dissolution of lipid domains over time.

**Figure 3:**
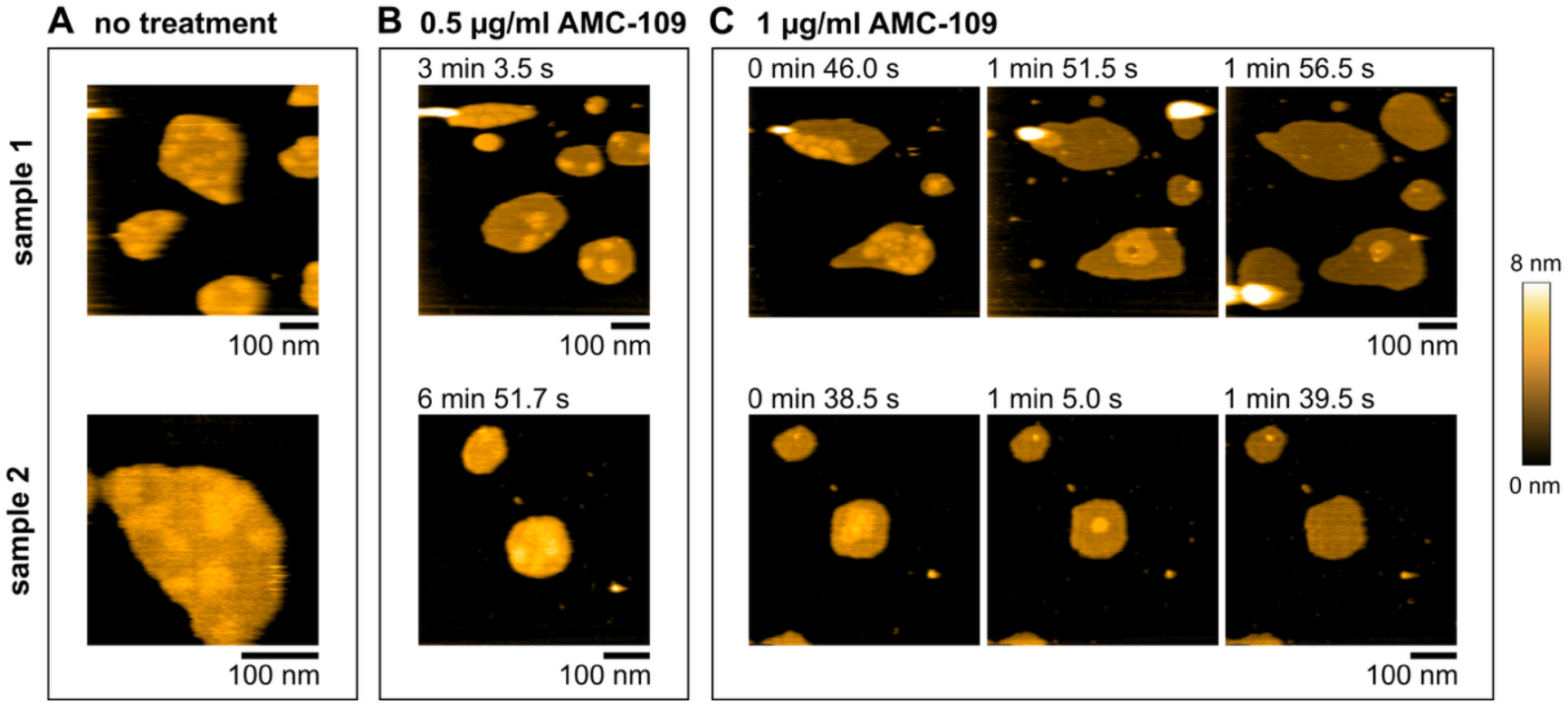
High-speed AFM images of MRSA lipid membranes upon exposure to sub-inhibitory concentrations of AMC-109. Separate MRSA lipid membrane patches (golden) lay flat on top of the mica support (black). 2 samples out of 4 measurements that yielded the same results are shown. (A) Without treatment, the membrane contains roughly circular domains, tens of nm in diameter (bright golden). Due to their fast movement, the domains are visible only with an image acquisition speed of 2–3 frames per second or faster (here 300–500 ms per frame). (B) At 0.5 µg/ml AMC-109, the domains start clustering and slow down their movement. (C) At 1 µg/ml AMC-109, some of the clustered domains dissolve. Three frames for this concentration at both samples illustrate this process in time.

We observed in the MRSA lipid membranes the domain clustering at 0.5 µg/ml and partial domain dissolution at 1 µg/ml AMC-109 in 4 separate samples. The results were further confirmed with standard AFM, where we observed untreated membranes of a uniform height, which subsequently formed clustered and immobile domains at 0.5 µg/ml. The acquisition of a single AFM image with our standard AFM setup took 10 minutes, meaning that only fully immobilized domains could be observed (Figure S5). We made the same observation on MSSA lipid extracts using a different standard AFM instrument with a time resolution of 2.5 minutes per frame. Here, we were able to visualize blurred domains in untreated MSSA lipid membranes, immobile domains after exposure to 0.5 µg/ml AMC-109, and partial domain dissolution at 1 µg/ml AMC-109 (Figure S6). The existence of the nanoscopic domains that we reported earlier^3,4^ is, hence, not bound to lipid membranes extracted from a single *S. aureus* strain, but can be observed across multiple strains including the methicillin-resistant one. Their shape, size, and fast movement that does not allow for tracking even with an imaging speed of 300 ms per frame is also strikingly similar in both MRSA (here) and MSSA (here and ref. ^3,4^) lipid membranes.

Moreover, the effects of AMC-109 on both MRSA and MSSA membranes are identical, affecting the membrane domains at the very same concentrations — importantly, at the same concentrations where potentiation of β-lactam antibiotics was observed (Figure 1–2). Hence, we can directly correlate the β-lactam resistance reversal in MRSA and the substantially increased β-lactam susceptibility of MSSA cultures with the membrane domain dissolution observed by AFM. The clustering of the domains that occurs at 0.5 µg/ml AMC-109 appears to be insufficient to fully boost β-lactam activity as evidenced by checkerboard assays (Figure 1). At 1 µg/ml, the threshold for domain dissolution is reached and, correspondingly, β-lactams activity is potentiated. This correlation supports our hypothesis that PBP2a, responsible for the resistance to methicillin and oxacillin, does not function in the absence of membrane domains, and aligns with the previous studies on reversing β-lactam resistance by disabling FMMs in MRSA by genetic modifications^16^ and phototherapy^13^. For the increased β-lactam activity in MSSA upon domain dissolution, we hypothesize that MSSA uses other types of PBPs located in the same domains that are rendered non-functional after domain dissolution. Likely, the activity, secretion, and/or recruitment of penicillinases in both MRSA and MSSA are among additional factors that are impaired. Thus, AMC-109 is paving the way for unobstructed activity of β-lactams. Together, AMC-109 and β-lactams coordinate a two-pronged attack stopping vital functions related to the structured and organized membrane (stopped by AMC-109) and the synthesis of the peptidoglycan cell wall (stopped by β-lactams). As the time-kill assays show, the MRSA culture could partially recover from the partial dissolution of the domains after exposure to 1 µg/ml AMC-109. Such recovery is abolished by the simultaneous inhibition of cell wall synthesis.

### AMC-109 disturbs membrane organization in living MRSA and MSSA and allows for oxacillin attack

We employed fluorescence microscopy to assess the effects of AMC-109 and oxacillin, singly and in combination, on live MRSA and MSSA cells. To this end, we used fluorescence dyes that specifically visualize (i) phase-separated membrane microdomains (DiIC12, Figure 4A and S7; Nile red, Figure S8–S9) ^35,36^, (ii) PBPs (Bocillin, Figure S10), (iii) the peptidoglycan precursor lipid II (Van-FL, Figure S8–S9), and (iv) membrane fluidity^30,32^ (Laurdan, Figure 4B–C, S11–12) ^35,37^. For single antibiotic treatment, the MIC was used to observe the changes in living bacterial cells. For combined treatment of MRSA, the same concentrations as in the time-kill assays (Figure 2) were used, i.e., 1 µg/ml AMC-109 + 0.25 µg/ml oxacillin. For combination treatment of MSSA, 1 µg/ml AMC-109 + 0.0156 µg/ml oxacillin were chosen based on the checkerboard assay results of this strain (Figure 1).

**Figure 4:**
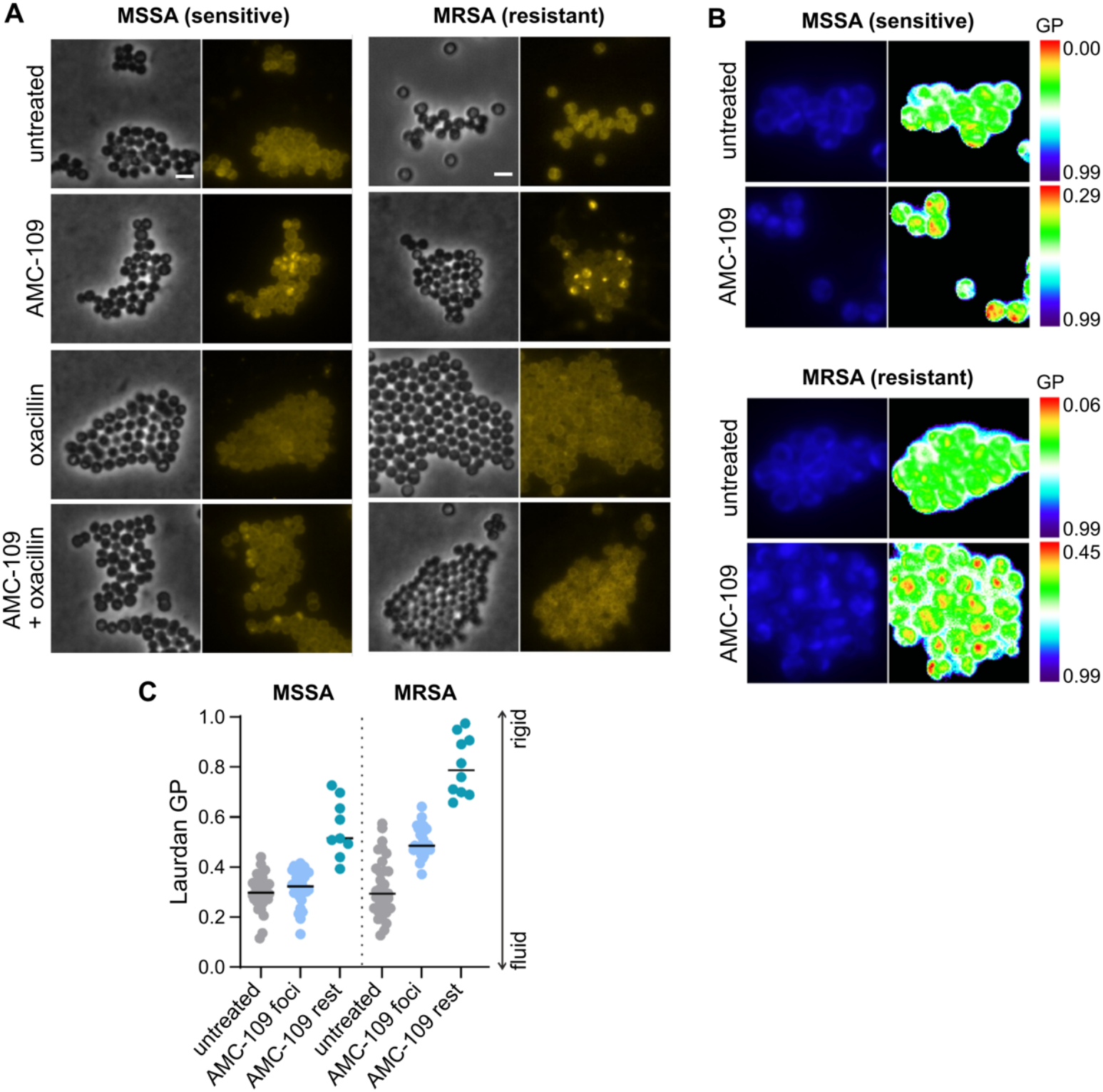
Fluorescence microscopy of MSSA and MRSA. (A) Phase contrast and fluorescence microscopy images of DiIC12-stained MSSA and MRSA. DiIC12 preferentially inserts into more fluid membrane domains due to its short hydrocarbon tail^35,38^. Bright yellow patches indicate fluid membrane domains in the bacterial membrane. Images were taken after 60 min of antibiotic treatment. Scale bars depict 2 µm. (B) Laurdan microscopy images of MSSA and MRSA showing Laurdan fluorescence at 350 nm excitation and 500 nm emission in blue and Laurdan GP maps calculated based on emission at 460 and 500 nm in rainbow LUT. A higher GP points to more rigid membranes. Scale bar depicts 2 µm. (C) Quantification of Laurdan GP measured from individual domains (foci) and out-of-domain (rest) regions after 60 min of antibiotic exposure.

The fluorescence microscopy data show that both MRSA and MSSA strains display a similar morphology typical for *S. aureus*. No prominent phase-separated domains were visible in the cell membranes of untreated MRSA and MSSA as evidenced by the fluidity-sensitive DilC12 (Figure 4A, S7) and Nile red (Figure S8–S9) dyes. This was expected as the nanometer-sized domains visible in extracted membranes under the high-speed AFM^3,4^ are below the spatial resolution of fluorescence microscopy, and is consistent with previous results on *S. aureus* cells^39^. AMC-109 treatment leads to clear phase separation into immobile, microscopically visible domains (Figure 4A, S7–9), which is in accordance with the high-speed AFM data (Figure 3). These phase-separated domains appear to be more fluid than the rest of the surrounding membrane as evidenced by accumulation of DilC12 (Figure 4A, S7) and Nile red (Figure S8–S9), which preferably partition into more fluid domains, whereby DiIC12 displays higher fluidity sensitivity than Nile red^36^.

To gain quantitative insight into the fluidity of these domains, Laurdan microscopy was performed. Laurdan binds to bacterial cell membranes and shifts its fluorescence intensity between two emission peaks at 460 and 500 nm, depending on the mobility of the surrounding lipid tails and hydration of the membrane. Thus, Laurdan generalized polarization (GP) is indicative of lipid spreading and gives a measure of membrane fluidity, a higher GP signifying a more rigid membrane environment^35,37^. Laurdan-stained MRSA and MSSA cells (Figure 4B, S11) showed clear phase-separated domains with an accumulation of laurdan in distinct foci. Laurdan GP quantification revealed that these domains have a higher fluidity than the surrounding rest membrane (yellow to red on the rainbow LUT scale), which is in line with DiIC12 and Nile red stainings. While these domains are indeed more fluid than the surrounding rest membrane (Figure 4C, S11 foci), they are of similar or slightly higher rigidity than untreated membranes. The rest membrane of treated cells is thus strongly rigidified (Figure 4C, S11 rest). This observation does not only indicate phase separation but also a general membrane rigidification effect of AMC-109. This notion was further confirmed in spectroscopic batch measurements (Figure S12). Overall reduction of membrane fluidity is in accordance with our previous AFM indentation experiments of MSSA lipid membranes, which showed an increased Young’s modulus after partial dissolution of the nanoscopic domains, likewise signifying membrane rigidification^3^.

Staining of MRSA and MSSA with Van-FL showed that the localization of the peptidoglycan precursor lipid II remained largely undisturbed by AMC-109 (Figure S8–S9). However, bocillin staining revealed displaced PBPs, a phenotype that coincided with the presence of immobile membrane foci Figure S10). This observation matches the literature on FMMs proposing that PBPs are localized in and depend on these domains^13,15,16,40,41^. The PBP localization in the domains, hence, supports our hypothesis that the total dissolution by membrane-domain-dissolving antibiotics is a valid target for enhancement of β-lactam activity.

Oxacillin alone does not influence membrane organization or fluidity (Figure 4A, S7, S12), but leads to strongly reduced bocillin staining, due to competition of oxacillin and bocillin (Figure S10). Oxacillin alone also leads to an aberrant Van-FL staining with pronounced septal accumulation of lipid II, which indicates impairment of peptidoglycan precursor incorporation into the cell wall and is in line with previous observations^42^ (Figure S8–S9). In cells treated with oxacillin and AMC-109, the bocillin phenotype was similar (Figure S10), while Van-FL staining did not show septal accumulation of lipid II (Figure S8–S9). This observation may be explained by an added effect of AMC-109 on lipid II synthesis due to energy limitation resulting from impairment of membrane function, a common downstream effect of membrane-targeting compounds^39,43,44^.

Interestingly, when AMC-109 was combined with oxacillin, the formation of immobile microscopic domains was considerably reduced in MSSA and abolished in MRSA (Figure 4A, S7). However, we still observed an overall membrane rigidification, which was particularly pronounced in MRSA (Figure S12, statistical significance of MRSA membrane rigidification by 1 µg/ml AMC-109 and by 1 µg/ml AMC-109 + 0.25 µg/ml oxacillin combination is p<0.01 and p<0.001, respectively). Taking into account the AFM data, showing domain dissolution following the formation of immobile foci (Figure 3), these observations suggest that domains are already fully dissolved at the timepoint of cell examination, a stark difference to single treatment with AMC-109, where immobile domains appeared to be stable in cells for at least 60 min (Figure 4A, S7). Oxacillin thus appears to enhance and/or accelerate the membrane-domain-dissolving ability of AMC-109, which could explain the prevention of CFU recovery upon combination treatment observed in Figure 2.

All our data combined, hence, support the hypothesis that dissolution of membrane domains by AMC-109 is the reason for the observed increase of β-lactam sensitivity in both MRSA and MSSA. Although some pore-forming peptides have been reported to synergize with β-lactams ^45–48^, they have not shown the potency to fully reverse the resistance in MRSA. With their undeniable potential to fully reverse the resistance in MRSA to multiple β-lactams, membrane-domain-dissolving antibiotics like AMC-109, hence, stand out as particularly powerful drug candidates to treat difficult-to-treat MRSA infections.

## CONCLUSION

The present study successfully demonstrates the hypothesis that membrane-domain-dissolving antibiotics reverse β-lactam resistance in MRSA through interference with membrane domain organization. We show that sub-inhibitory concentrations of the membrane-domain-dissolving antibiotic AMC-109 lower the MICs of three β-lactam antibiotics (penicillin G, oxacillin, methicillin) 2–256 fold in both MRSA and MSSA. Importantly, the oxacillin MIC against MRSA is reduced below the clinical resistance breakpoint defined by the FDA and EUCAST, effectively reversing resistance. Based on our results, we propose the combination of oxacillin with a membrane-domain-dissolving agent such as AMC-109 to restore activity against MRSA. This new combination appears to be highly effective, achieving a reduction from 10^7^ to 10^3^ CFU/ml of MRSA within 4 hours.

We could pinpoint the mechanism behind this resensitization effect to interference of AMC-109 with membrane domains, resulting in disturbance of PBP2a localization. Thus, concentrations of the membrane-domain-dissolving antibiotic achieving β-lactam potentiation cluster and subsequently partially dissolve membrane domains in MRSA lipid extracts. Similarly, the same concentrations fuse membrane domains in living bacteria into immobile foci and disturb PBP localization. Hence, we were able to link restored β-lactam susceptibility to domain clustering and dissolution in both living bacterial cells and membrane extracts from the same strains. Furthermore, fluorescence microscopy of living bacteria suggests that combined treatment of membrane-domain-dissolving antibiotic with β-lactam enhances and/or accelerates the domain dissolution effect, which in turn allows for the full susceptibility renewal of MRSA to β-lactams.

For MRSA, we can explain restored β-lactam susceptibility with the dissolution of membrane domains that constitute the site of PBP2a oligomerization (Figure 5). It is, however, likely that additional mechanisms are at play. The improved susceptibility of MSSA to all tested β-lactams upon combination with AMC-109 cannot be explained by a lack of PBP2a oligomerization as MSSA does not possess PBP2a. It is thus reasonable to assume that disruption of membrane domain organization by AMC-109 also affects other PBPs as well as other membrane-bound proteins such as penicillinases and efflux pumps. Such additional effects of membrane-domain-dissolving antibiotics on membrane protein function as well as accelerated domain dissolution by the oxacillin/AMC-109 combination remain to be investigated in further studies.

**Figure 5:**
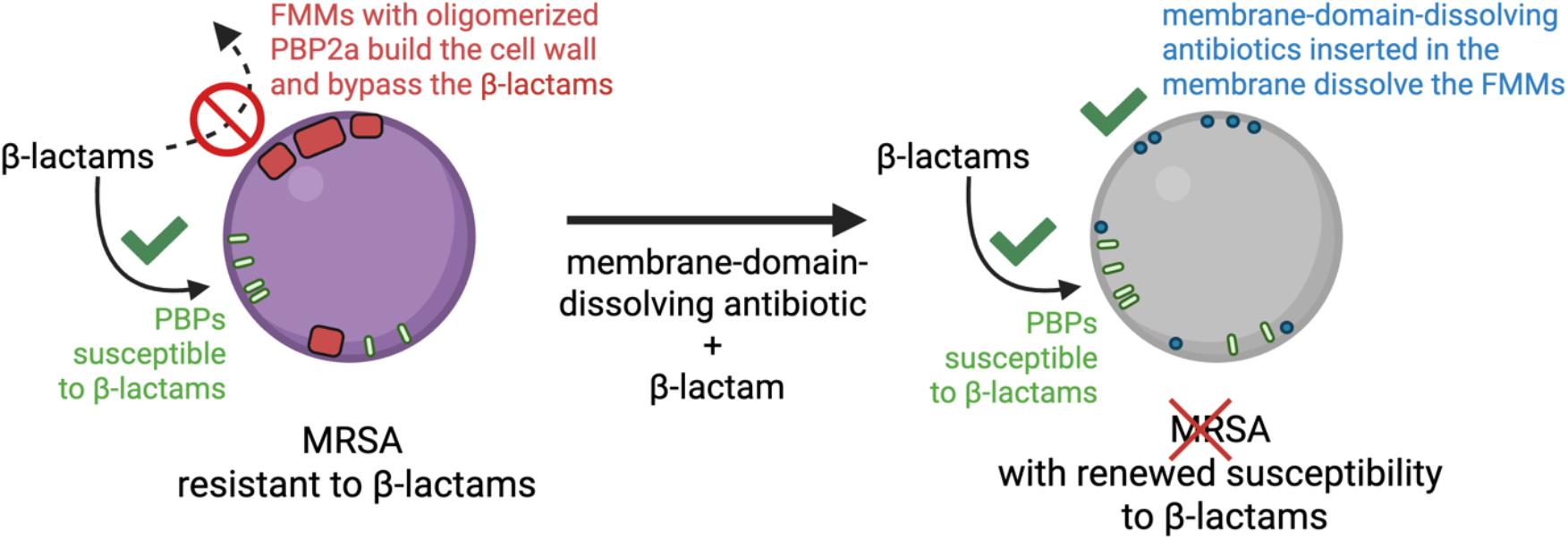
Model of β-lactam susceptibility renewal in MRSA based on the data presented in this study.

In conclusion, AMC-109, with its ability to fully restore the susceptibility of MRSA to multiple β-lactams below their clinical breakpoints, stands out as particularly powerful drug candidate to ‘treat the untreatable’ in the fight against antimicrobial resistance.

## MATERIALS AND METHODS

### Materials

AMC-109 (previously LTX-109^31^) was obtained as a dry powder from Amicoat AS, Norway and stored at -20°C. Oxacillin sodium monohydrate and methicillin sodium salt were purchased as dry powders from MedChemExpress and stored at 4°C. Penicillin G sodium salt was purchased as a dry powder from Sigma Aldrich and stored at 4°C. Mueller Hinton Broth (MHB) for bacteria cultivation was purchased from Sigma Aldrich and stored at room temperature. Fluorescent dyes Nile red, Van-FL, Bocillin, and DiIC12 were purchased from Invitrogen, Laurdan from Anaspec and stored at -20 °C.

All the antibiotic powders were dissolved in dimethyl sulfoxide (DMSO) to 10 mg/ml stock solutions and stored at -20°C. The DMSO stock solutions were used for a maximum of 6 months in case of AMC-109, and for the maximum of 1 month in case of oxacillin, methicillin, and penicillin G. For the experiments, DMSO stock solutions of antibiotics were diluted in MHB, water, or buffer to the desired concentration and used the same day.

### Bacterial strains

Clinical isolate methicillin-resistant *Staphylococcus aureus* (MRSA) and methicillin-susceptible *S. aureus* (MSSA) strain ATCC 29213 were acquired from Dr. E. Bathoorn from the University Medical Center Groningen. These strains were used for minimum inhibitory and bactericidal concentration testing (MIC and MBC), checkerboard assays, time-kill assays, and fluorescence microscopy imaging. MRSA was also used for lipid extraction and subsequent AFM imaging.

A second used MSSA strain RN4220, a restriction-deficient and prophage-cured NCTC8325-4 derivative, was a gift from prof. J. M. van Dijl from the University Medical Centrum Groningen. Lipid extracts from this MSSA strain were used for AFM imaging.

### Cultivation of bacteria for activity experiments

All bacterial culture stocks were stored at -80°C in 7% DMSO. The bacterial stocks were streaked onto a blood agar plate (5% sheep blood) and cultured overnight at 37°C. Blood agar plates with bacteria were stored at 4°C and used for a maximum of 1 month. A single colony of bacteria from the blood agar plate was transferred to 10 ml of sterile MHB medium to prepare a preculture, which was incubated for 24 h at 37°C in static condition. Hereafter, 2.5 ml of the preculture was transferred into 50 ml of sterile MHB in a 100 ml Erlenmeyer flask, which was incubated at 37°C overnight (16–18 h) while shaking at 150 rpm. The following day, the main culture was transferred to an ultracentrifuge tube and centrifuged at 5000 g at 10°C for 5 min to separate a pellet of the bacteria cells from the medium. The supernatant was discarded and the pellet was washed and resuspended in 10 ml PBS. The centrifuging and washing steps were repeated three times. Hereafter, the pellet was resuspended in 5 ml of sterile PBS, into which a tip sonicator (Vibra cell by Sonics and Materials Inc.) was submerged. The bacteria were tip sonicated for 30 s with sonication pulses of 1 s duration with 2 s pause in between the pulses with an amplitude of 60%. Then the density of the separated individual bacteria per ml was counted. For the bacteria counting, the sonicated bacteria culture in PBS was diluted to achieve a density of roughly 10 bacteria cells in one square well of the Burker Turk counting chamber. From the bacteria counting, the density in the main stock was calculated, and the main stock was diluted for the desired starting bacteria/ml for a given experiment.

### Minimum inhibitory concentration (MIC) and minimum bactericidal concentration (MBC) testing

96-well plates with a round bottom were used for MIC testing. 200 µl of MHB growth medium with double of the desired maximum antibiotic concentration was added to the first well of each row. This doubly concentrated antibiotic was diluted twice with 100 µl of MHB in the following well in the row, mixed well, and subsequently diluted to the next well until a gradient of two-fold diluted concentrations was created in the row with 100 µl of MHB with antibiotic in each well. The last well of each row was left without any antibiotic addition as a growth control. 100 µl of 2.10^5^ bacteria/ml in MHB were added to each well achieving 1.10^5^ bacteria/ml in each well and the correct concentration of antibiotics. The 96-well plate was then sealed with parafilm with a small gap for air exchange and incubated at 37°C for 24 hours in static condition. Afterward, the MIC was determined as the first antibiotic concentration with no visible bacterial growth. The MBC was determined from the wells with no bacteria growth, and from the highest antibiotic concentration with growth to have a positive growth control. The suspension in each well was mixed with a pipette and a 10 µl drop was deposited on an MH agar plate and allowed to dry. The MBC was determined as the lowest antibiotic concentration with no bacteria growth on the agar plate after 24 hours of incubation at 37°C. After taking out the 10 µl drops, the MIC 96-well plate was returned to 37°C and incubated for another 24 hours to control the MIC after 48 hours of antibiotic activity. MIC and MBC of all antibiotics against MRSA and MSSA were evaluated in duplicates in 6–7 independent biological replicates, yielding similar results. Table 1 shows the highest MIC and MBC values from these repeated measurements. MIC at 48 hours was always the same or one dilution higher than the MIC at 24 hours. MIC at 48 hours usually coincided with the MBC value (Table S1).

### Checkerboard assays

To prepare the 96-well plates with round bottom for the checkerboard assays, the concentration gradients of the first antibiotic in columns and of the second antibiotic in rows need to be achieved. Helping 96-well plates were used to prepare both gradients separately. In the helping plates, the gradient in columns with the quadruple total concentration of the first antibiotic, and the gradient in rows with the quadruple total concentration of the second antibiotic were prepared. In the final experimental plate, we combined (i) 50 µl of each well from the first helping plate, (ii) 50 µl of each well from the second helping plate, and (iii) 100 µl of 2.10^5^ bacteria/ml in MHB to achieve correct concentrations of all components. The plates were sealed with parafilm leaving a small gap for air exchange and incubated for 24 hours at 37°C. After 24 hours, the MIC of the combination treatments was determined. Also here, MBC was tested in all wells with no growth and in the wells with the highest concentration of β-lactams with growth. MBC and MIC at 48 hours were tested the same way as described above. Checkerboard assays were performed in 4–5 independent experimental days, each time with new bacteria cultivation. The concentration of AMC-109 which increases the β-lactam activity was the same in all repeated experiments. 0.5 µg/ml AMC-109 leads to slight improvement, 1 µg/ml AMC-109 yields to multiple times improvement in MIC. The level of MIC reduction of β-lactams varied between the repetitions (4x–512x MIC reduction).

### Time-kill assays

Four 250 ml Erlenmeyer flasks were prepared with 100 ml of 1.10^7^ bacteria/ml in MHB. The following treatments were added to the flasks: (i) no treatment, (ii) 1 µg/ml AMC-109 + 0.25 µg/ml oxacillin, (iii) 1 µg/ml AMC-109, (iv) 0.25 µg/ml oxacillin. Ten times 10 ml were aliquoted from each flask into sterile glass tubes, leading to 10 tubes with each treatment. The tubes were incubated at 37°C under static conditions. At each time point, one tube per treatment condition was taken out of the incubator. The tubes were checked for optical density (OD 600, Figure S3) with the spectrometer Genesys 30 from Thermo Scientific™. When the OD was above 0.6, the sample was diluted to a value below 0.6. Each sample was then diluted 10-fold (180 µl + 20 µl) in PBS using a 96-well plate, and then diluted 10 times into the next wells achieving dilutions 10^-1^ to 10^-8^ times. Three 10 µl droplets of each dilution were placed on the MH agar plate and allowed to dry. 100 µl of the undiluted suspension were transferred on another MH agar plate and spread over the plate with a spatula. The plates from the measurement time points 0–7 hours were collected at room temperature and then incubated 16–18 hours at 37°C. The next day, colony forming units (CFU) were counted at the dilutions where clear separated colonies were formed. The reported CFU is the average of all CFU counts from the three 10 µl droplets of all concentrations, where the counting was possible. The procedure was repeated for the time points 24 and 48 hours on the following days. Overall, the time points tested were 0, 0.5, 1, 2, 3, 4, 5, 6, 7, 24, and 48 hours. The experiment with all treatment conditions was repeated in 3 biological replicates.

### Lipid extraction

Lipids were extracted following the modified Bligh & Dyer extraction protocol^49^ as we reported earlier^3^. For *S. aureus* RN4220, NCTC8325-4 derivative (MSSA), the same stock of the lipid extract as in ^3^ was used. New extraction was done from the clinical isolate of MRSA. One bacteria colony from blood agar plate was inoculated into 10 ml of MHB and incubated for 24 hours at 37°C in static condition. Hereafter, 10 ml of the preculture was transferred into 200 ml of MHB for the main culture and incubated for 24 hours at 37°C while shaking with 150 rpm. The following day, the main culture was transferred into centrifuge tubes and centrifugated at 5000 g at 10°C for 5 min, the supernatant was discarded, and the pellet resuspended in 10 ml of demi water. The samples were pooled into a 50 ml centrifuge tube and centrifuged for 5 min at 10°C at 5000 g. The supernatant was discarded and the bacterial pellet was dried in air. The following day the pellet was dissolved in water/chloroform/methanol in ratios 0.8:1:2 and stirred at 4°C overnight to kill the bacteria and disassociate the lipids from the rest of the cellular material. The sample was centrifuged for 15 min at 900 g at 4°C. The collected supernatant contains remains of the original 0.8:1:2 water/chloroform/methanol suspension. The wet supernatant was added 1:1 into water/chloroform in a 1:1 ratio, so the final water/chloroform/methanol ratio of 0.9:1:1 was achieved. The solution was incubated for 2 days at room temperature for the phase separation. The bottom layer, which contains the lipids, was collected and dried at room temperature. The dry lipid extract was weighted, dissolved in the concentration 10 mg/ml in chloroform, and stored at -20°C until use.

### Atomic force microscopy

MSSA and MRSA lipid extracts in chloroform were pipetted into a glass vial. An argon stream was used to gently evaporate the chloroform, and the lipid film on the glass was left to fully dry in a vacuum for >1 hour. PBS was added to the lipid film in the concentration 0.3–1 mg/ml. The lipid suspension was then vortexed for ∼2 min, which was followed by 5–7 cycles of freeze and thaw using liquid nitrogen and a warm water bath. Multiple freeze and thaw cycles were required to fully detach all the lipid material from the glass. The resuspended liposomes were then stored in 4°C and used for the maximum of 3 days.

High-speed AFM imaging was performed using a RIBM (Japan) machine in amplitude modulation tapping mode in liquid^3,4,50,51^ at room temperature. Ultrashort cantilevers USC-F1.2-k0.15 (NanoWorld, Switzerland) with spring constants of 0.15 N/mm, a resonance frequency of ∼0.6 MHz, and a quality factor of ∼2 were used. The cantilever free amplitude was set to 1 nm, and the set-point amplitude for a surface detection around 0.8 nm was used. Images were taken with a frame rate of 300 ms–2 s per frame. A mica surface of 1.5 mm in diameter was glued on top of a 5 mm high glass rod which was used as the sample stage for the high-speed AFM. The glass rod was then attached to the scanner Z-piezo using a nail polish as glue. A 2–3 µl drop of 0.03–0.04 mg/ml MRSA lipid extract liposomes in PBS was deposited on freshly cleaved mica and left to adhere for >10 min. The buffer was washed out 5 times with a fresh drop of PBS to press the liposomes on the mica surface and force them to collapse into supported membrane patches. The scanner head was then put upside down into a small liquid chamber containing the cantilever and filled with 75 µl of PBS. Untreated membranes were first imaged, and then 5-15 µl of AMC-109 was added step by step to achieve the desired 0.5, 1, and 2 µg/ml final concentration in the sample solution.

AFM imaging of MSSA ATCC 29213 lipid extracts was performed using Bruker Nanoscope Catalyst AFM in peak force quantitative imaging mode using qp-BioAC cantilevers (NanoAndMore GmbH) with a nominal spring constant of 0.1 N/m (0.06–0.18 N/m). A sheet of mica was glued on top of a cover glass with the second glue and cleaved before each experiment. A circle with a PAP pen was drawn around the freshly cleaved mica to create a hydrophobic barrier. A drop of 10 µl of the 0.03 mg/ml liposome suspension was deposited into the hydrophobic circle on top of the cleaved mica and left to adhere for >10 min. The PBS buffer was washed out similarly to the high-speed AFM measurements to force the formation of supported membrane patches. A 50 µl droplet of the fresh PBS buffer was used for imaging. AFM imaging of lipid extracts from MSSA RN4220, NCTC8325-4 derivative, was performed with a JPK Nano Wizard Ultra Speed AFM using qp-BioAC cantilevers (NanoAndMore GmbH) with a nominal spring constant of 0.06 N/m (0.03–0.09 N/m). Here, the liquid imaging well was prepared by gluing a small piece of mica with epoxy glue on top of the microscopy glass slide and attaching a glass ring around it with biocompatible two-component glue (Bruker Nano GmbH). A 10 µl drop of 0.03 mg/ml liposomes in PBS was deposited on freshly cleaved mica, left to adhere for >10 min, and washed out. The liquid well was filled with PBS to a total volume of 1 ml for imaging. All the AFM imaging experiments were performed in PBS at room temperature. First, untreated membranes were imaged. Afterwards a droplet of concentrated antibiotic solution in PBS was added step by step to achieve the desired final concentration of 0.5, 1, and 2 µg/ml.

### Fluorescence microscopy and spectroscopy

Fluorescent dyes were prepared and stored as follows. Nile red was prepared as 1 mg/ml stock solution in sterile DMSO. DiIC12 was prepared as single-use aliquots with a concentration of 100 µg/ml in DMSO. Laurdan was prepared as 10 mM stock solution in DMF and diluted to a 1 mM working stock on the day of the experiment. Bocillin was prepared as 1 mg/ml stock in sterile water. Van-FL was prepared as 1 mg/ml single-use aliquots in sterile water and mixed 1:1 with unlabeled vancomycin immediately prior to use. All stock solutions were stored at -20 °C until further use.

*S. aureus* strains were grown in MHB at 37 °C under constant shaking (180 rpm). Overnight cultures were diluted to a starting OD_600_ of 0.05 and allowed to grow to an OD_600_ of 0.3 prior to addition of antibiotics. Antibiotic concentrations were chosen based on MIC and checkerboard data (Table 1, Figure 1). MSSA was treated with 4 µg/ml AMC-109, 0.5 µg/ml oxacillin, 1 µg/ml AMC-109 + 0.0156 µg/ml oxacillin, or left untreated as control. MRSA was treated with 2 µg/ml AMC-109, 3 µg/ml oxacillin, 1 µg/ml AMC-109 + 0.25 µg/ml oxacillin, or left untreated. Microscopy images were taken after 30 and 60 min of treatment.

Fluorescent stainings were performed as follows. All staining times were included in the final treatment times for keeping the examined timepoints consistent between experiments. For Nile red and bocillin co-staining, cell samples were stained with 1 µg/ml Nile red for 5 min and 1 µg/ml bocillin for 10 min. To avoid misinterpretation of possible fluorescence bleed-through artefacts, samples stained with each dye individually were observed in addition to the co-stained samples. No bleed-through artefacts were observed in these experiments. Van-FL staining was performed at a final Van-FL concentration of 1 µg/ml for 2 min. DiIC12 staining was performed as described previously^36^. In short, 1 µg/ml DiIC12 was added to the culture immediately after dilution to OD 0.05. A constant concentration of 1% DMSO was maintained to prevent precipitation of the dye. After reaching an OD_600_ of 0.3, cells were washed four times in pre-warmed MHB + 1% DMSO and resuspended in the same medium, followed by antibiotic treatment. For laurdan microscopy and spectroscopy^36^, cells were stained with 1 µM laurdan for 5 min. A constant concentration of 1% DMF was maintained to avoid precipitation of the dye. Cells were washed four times with pre-warmed laurdan buffer (PBS, 1% DMF, 0.2% glucose) and resuspended in the same buffer prior to antibiotic treatment.

Fluorescence microscopy was performed on a Nikon Eclipse Ti2 inverted fluorescence microscope equipped with a CFI Plan Apochromat phase contrast objective (DM Lambda 100X Oil N.A. 1.45, W.D. 0.13mm, Ph3), a Lumencor Sola SE II FISH 365 light source, a Photometrics PRIME BSI camera, an Okolab incubator, and Nis ELEMENTS AR 5.21.03 software. Filter settings (EX: excitation/bandwidth, DM: dichroic mirror, EM: emission/bandwidth) were as follows: EX 562/40, DM 593, EM 640/75 for Nile red, EX 466/40, DM 495, EX 520/50 for Van-FL and bocillin, EX 531/40, DM 562, 593/40 for DiIC12, and EX 377/50, DM 409, EM 447/60 and EM503/40 for laurdan. All microscopy images were processed and analyzed with Fiji. Laurdan microscopy was analyzed with the ImageJ ‘Calculate GP’ plugin^35,52^. Fluorescence spectroscopy was performed in a BMG Clariostar Plus platereader with a 350 nm excitation wavelength and 460 and 500 nm emission wavelengths. Laurdan GP was calculated according to the formula GP=(I_460_-I_500_)/(I_460_+I_500_)^36^.

### Statistical analysis

For statistical test of the significance of the observed changes 2-tail Student’s t-test comparing two data sets was used, assuming the same variance of the two normally distributed data sets in all cases when the standard deviations of the two compared data groups deviated by less than 2-times factor. T-test with unequal variance was used when the two compared data sets had a larger difference between their standard deviations. For data presented in Figure 2, log_10_(CFU/ml) in 24h, and 48h were compared to the value at 5h to show the significance of the recovered growth of bacteria culture treated with 1 µg/ml AMC-109. For data presented in Figure S12, Laurdan GP values after treatments were compared to the untreated sample values. Asterisks in Figures illustrate significance of the p-value resulting from the t-test, * indicates p<0.05, ** is used for p<0.01, and *** for p<0.001.

## ACKNOWLEDGEMENTS

We thank Arnold J.M. Driessen (University of Groningen) for the gift of *S. aureus* strain RN4220 and for support. We thank Janny de Wit (University of Groningen) for her invaluable help with *S. aureus* cultivation and lipid extraction. We thank Erik Bathoorn (University Medical Center Groningen) for providing the MRSA and MSSA (ATCC 29213) strains of *S. aureus*. AM thanks the Foundation to Prevent Antibiotic Resistance for support through the Research Grant 2023 and the Nederlandse organisatie voor Wetenschappelijk Onderzoek (NWO) for support through a Physics/f grant (no. 680-91-007) and an XS grant (OCENW.XS24.2.021). MW thanks Martin Andersson (Chalmers University of Technology) for access to BSL-II facilities.

## AUTHOR CONTRIBUTIONS

AM conceptualized and designed the work. AM and WW performed, analysed, and interpreted the MIC, MBC, checkerboard and time-kill assays. AM performed, analysed and interpreted the high-speed and standard Bruker Catalyst AFM experiments. MG performed the standard JPK Nano Wizard Ultra Speed AFM experiments under the supervision by AM and WHR. WHR surveyed the standard JPK Nano Wizard Ultra Speed and high-speed AFM experiments. MW performed, analysed, and interpreted the fluorescence microscopy and spectroscopy experiments. WS and JSMS consulted the activity of AMC-109 and provided technical support and AMC-109 material for this study. RS supervised the work. AM wrote the article. RS, WW, WHR and MW revised the written version of the article. All the authors have approved the final version to be published.

## DATA AVAILABILITY STATEMENT

Data that support findings of this work can be found in the manuscript and its Supplementary Information. Raw AFM images, individual frames for the high-speed AFM observations, and raw fluorescence microscopy images and data from experimental repetitions are available upon request.

