## Supplementary Information for "Treating the untreatable: Reversing β-lactam resistance in MRSA by membrane-domain-dissolving antibiotics"

### Supporting Information for
### Treating the untreatable: Reversing β-lactams resistance in MRSA by membrane-domain-dissolving antibiotics

Adéla Melcrová[1*], Willem Woudstra[1], Michaela Wenzel[2,3], Mariella Gabler[4], Wenche Stensen[5], John S. M. Svendsen[5], Wouter H. Roos[4], Romana Schirhagl[1]

[1] Biomaterials and Biomedical Technology, University Medical Center Groningen, The Netherlands
[2] Division of Chemical Biology, Department of Life Sciences, Chalmers University of Technology, Gothenburg, Sweden
[3] Center for Antibiotic Resistance Research in Gothenburg (CARe), Gothenburg, Sweden
[4] Molecular Biophysics, Zernike Institute for Advanced Materials, University Groningen, The Netherlands
[5] Department of Chemistry, UiT Arctic University of Norway, Norway

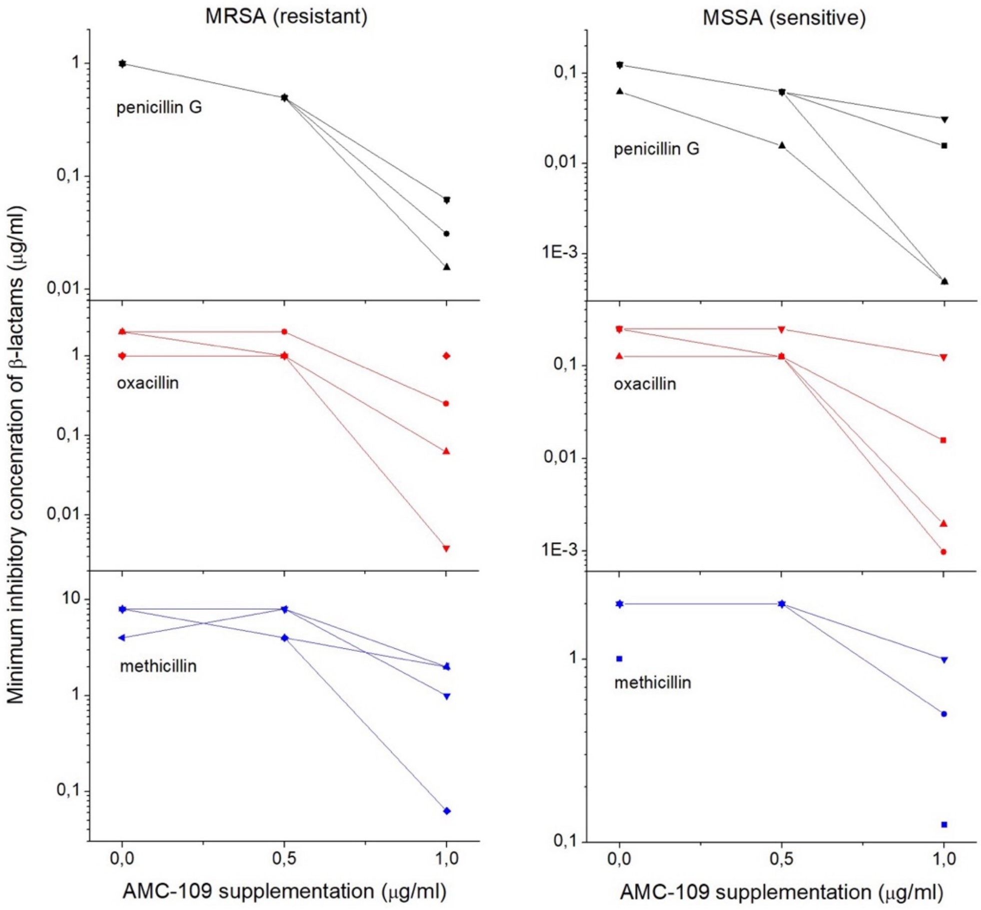


**Figure S1**: Minimum inhibitory concentrations (MICs) of β-lactams drop upon supplementation with AMC-109 against both MRSA and MSSA. Graphs show data points from 3–4 replicate experiments. For penicillin G, the minimum observed MIC lowering at 1 µg/ml AMC-109 supplementation is 8-fold ($2^3$) and the maximum MIC lowering is 256-fold ($2^8$; shown in Figure 1 MSSA penicillin G/AMC-109). For oxacillin, the minimum and maximum MIC lowering varies between 4-fold and 256-fold. For methicillin, between 2-fold and 256-fold. Connecting lines are missing in cases where the middle point (0.5 µg/ml) was non-conclusive due to contamination of the 96-well plate.

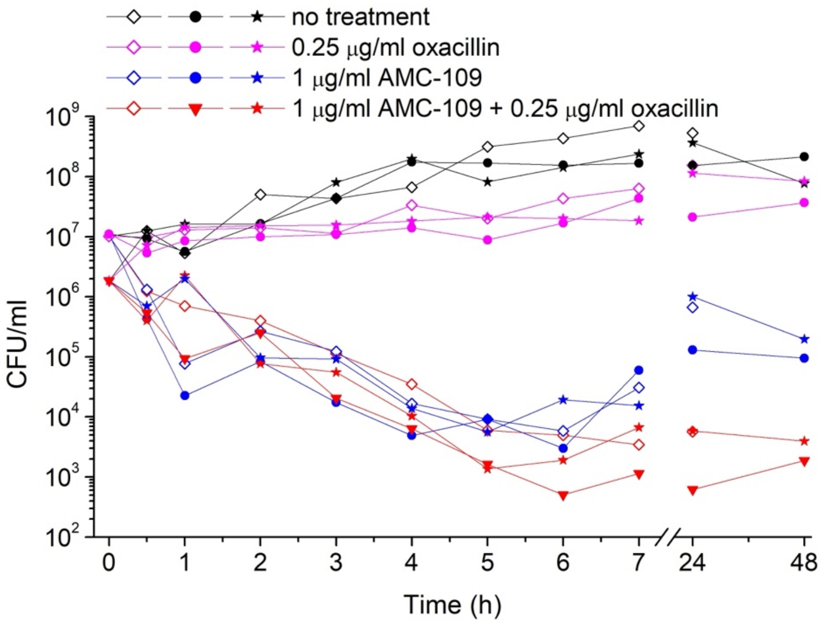


**Figure S2**: Time-kill assays of MRSA upon the exposure to no treatment (black), 0.25 µg/ml oxacillin (magenta), 1 µg/ml AMC-109 (blue), and the combination therapy of 1 µg/ml AMC-109 + 0.25 µg/ml oxacillin (red)). Data points are individual values gained from 3 biological replicates. 48 h data were gathered from 2 biological replicates.

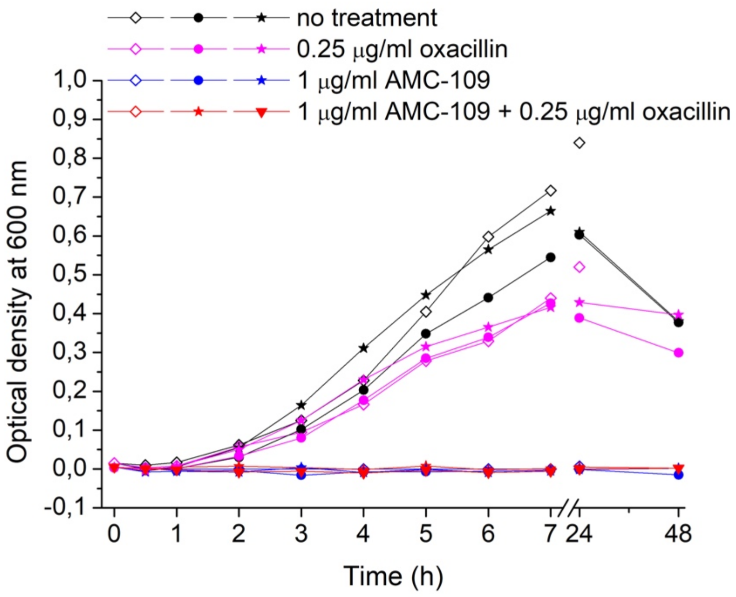


**Figure S3**: Optical density of MRSA cultures corresponding to the same cultures shown in Figure S2 upon the exposure to no treatment (black), 0.25 µg/ml oxacillin (magenta), 1 µg/ml AMC-109 (blue), and the combination therapy of 1 µg/ml AMC-109 + 0.25 µg/ml oxacillin (red). Data points are individual values gained from 3 biological replicates. 48 h data were gathered from 2 biological replicates.

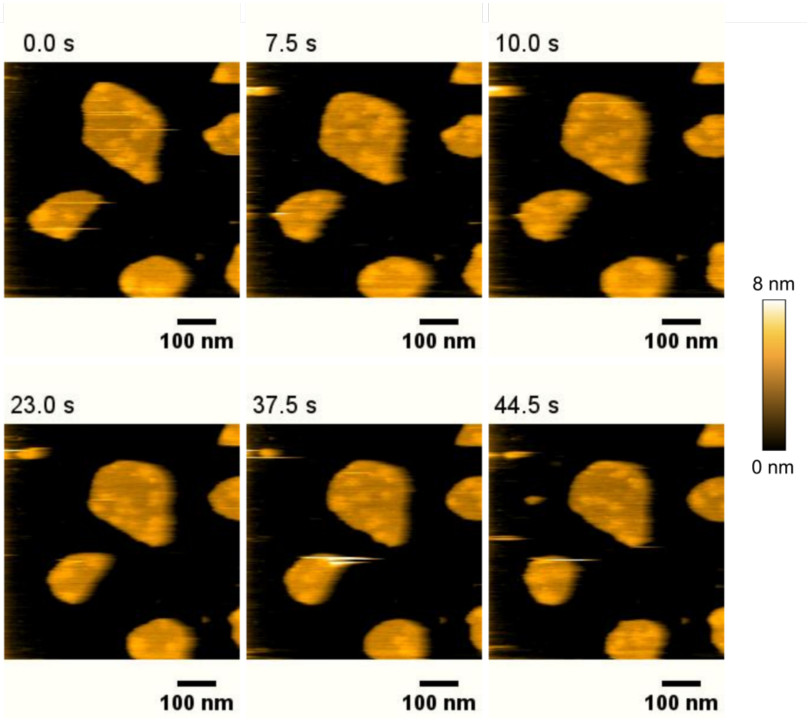


**Figure S4:** Snapshots from the high-speed AFM Supporting video 1 showing the movement of membrane domains in lipid membranes extracted from MRSA. The membrane patches (golden) are supported on mica (black). Separated domains (bright golden) are highly mobile. High-speed AFM video was acquired with an acquisition speed of 2 frames per second.

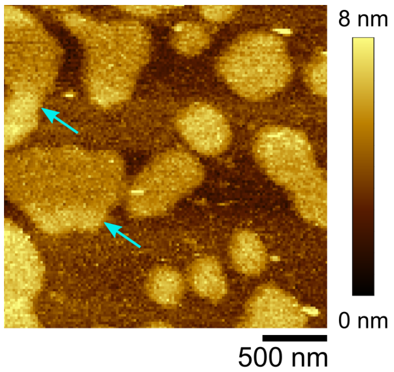


**Figure S5:** AFM image of MRSA lipid membranes exposed to 0.5 µg/ml AMC-109. Separate membrane patches (golden) are supported on mica (brown). Blue arrows show higher parts of the membrane (bright golden), which indicate the presence of stopped and clustered domains. Measured on Bruker Catalyst AFM with an image acquisition time of 10 min.

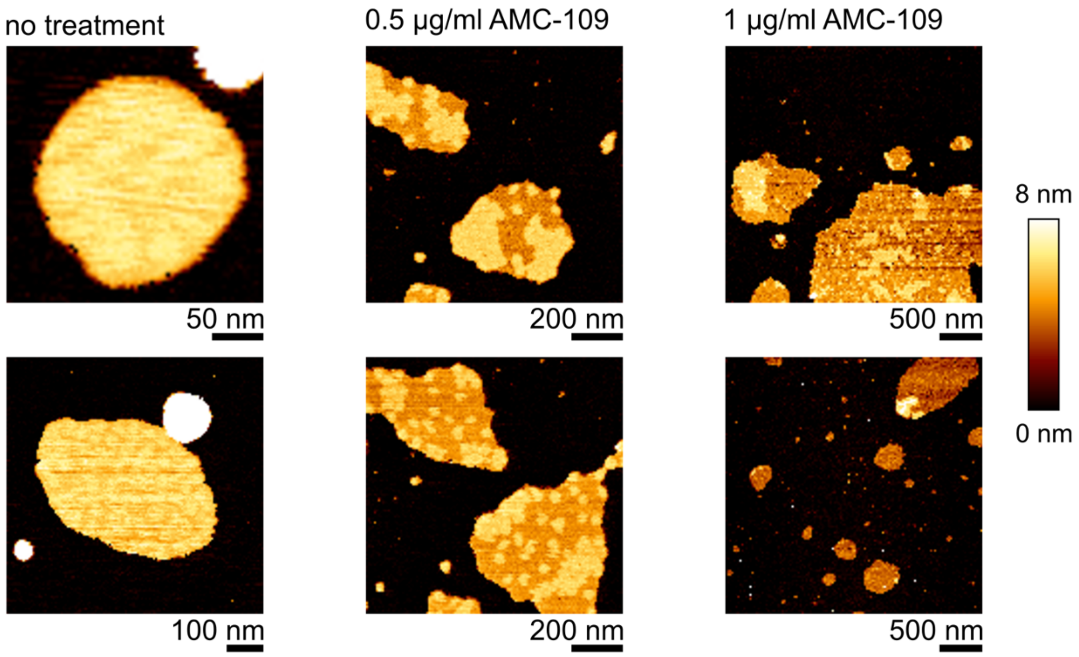


**Figure S6:** AFM images of MSSA lipid membranes exposed to 0, 0.5, and 1 µg/ml AMC-109. Separate membrane patches (orange/yellow) are supported on mica (black). Higher parts of the membrane are visible as bright yellow. Two representative images are shown for each condition. At no treatment, individual membrane domains are blurred together. After exposure to 0.5 µg/ml AMC-109, the domains stop their movement and can be visualized with the standard AFM at an image acquisition time of 2.5–10 min. At 1 µg/ml AMC-109, some of the domains are dissolved into the surrounding membrane. Images were recorded with a JPK Nano Wizard Ultra Speed AFM.

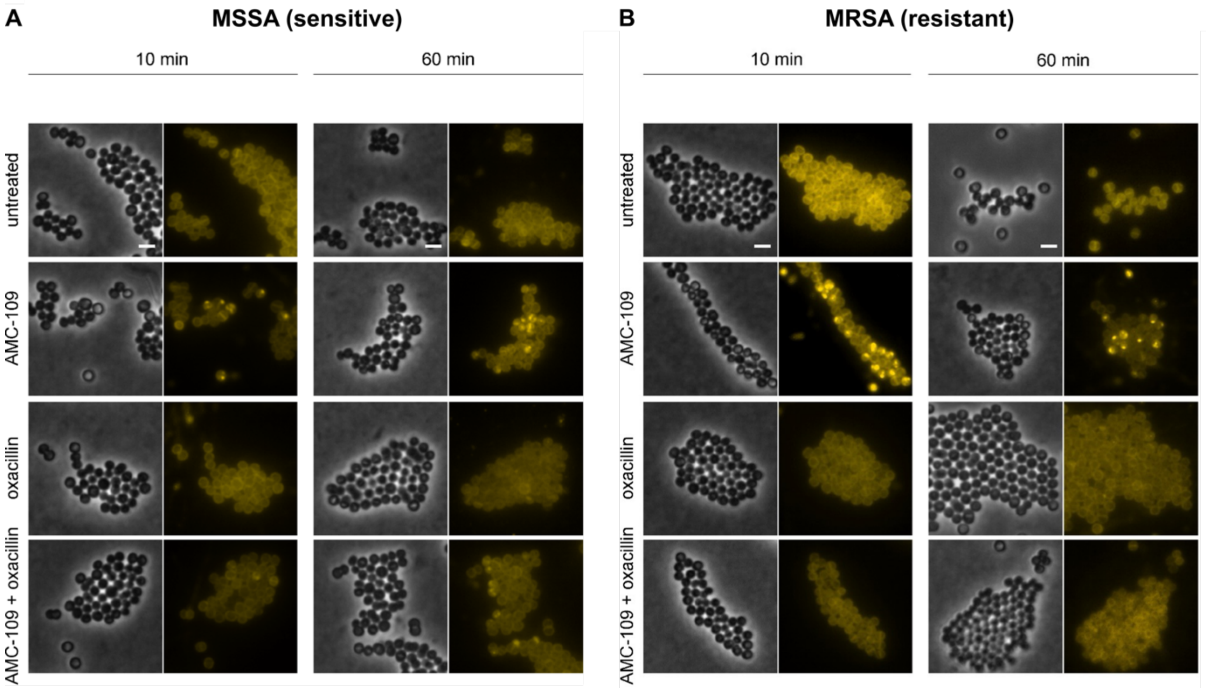

**Figure S7**: Phase contrast and fluorescence microscopy images of DiIC12-stained (A) MSSA and (B) MRSA taken after 10 and 60 minutes of antibiotic treatment. DiIC12 preferentially inserts into more fluid membrane domains due to its short hydrocarbon tail[34,37]. Bright yellow patches indicate fluid membrane domains in the bacterial membrane. Scale bars depict 2 µm.

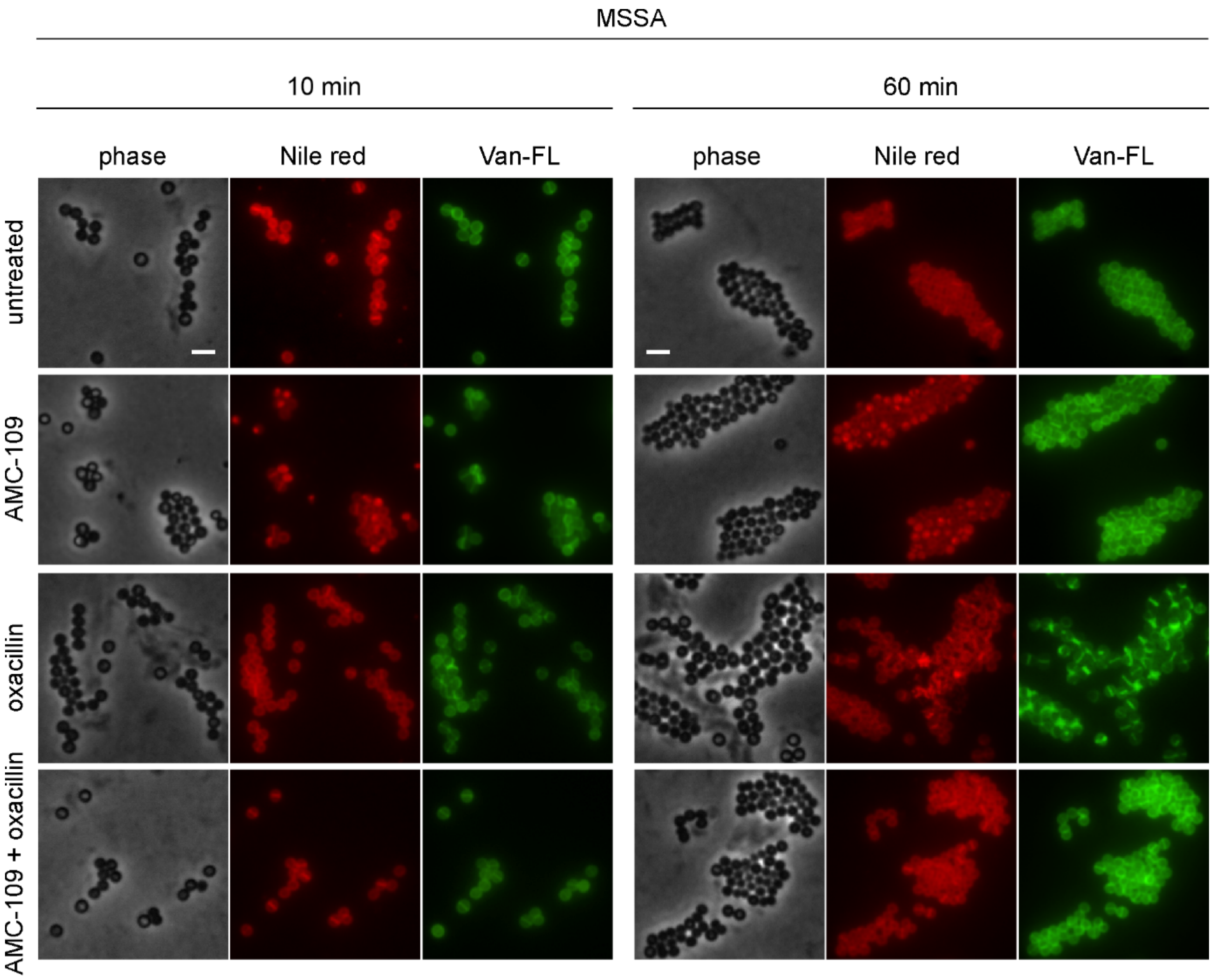


**Figure S8:** Microscopy images of phase contrast, Nile red, and Van-FL staining of MSSA. Nile red is a membrane dye that unselectively stains bacterial cell membranes but partitions into the fluid phase upon phase separation. Accumulation of Nile red in bright spots indicates fluid membrane domains. Van-FL is a fluorescently labeled vancomycin derivative that binds to lipid II, indicating its localization and serving as proxy for sites of active cell wall synthesis. MSSA was treated with 4 µg/ml AMC-109, 0.5 µg/ml oxacillin, 1 µg/ml AMC-109 + 0.0156 µg/ml oxacillin, or left untreated as control. Scale bars represent 2 µm.

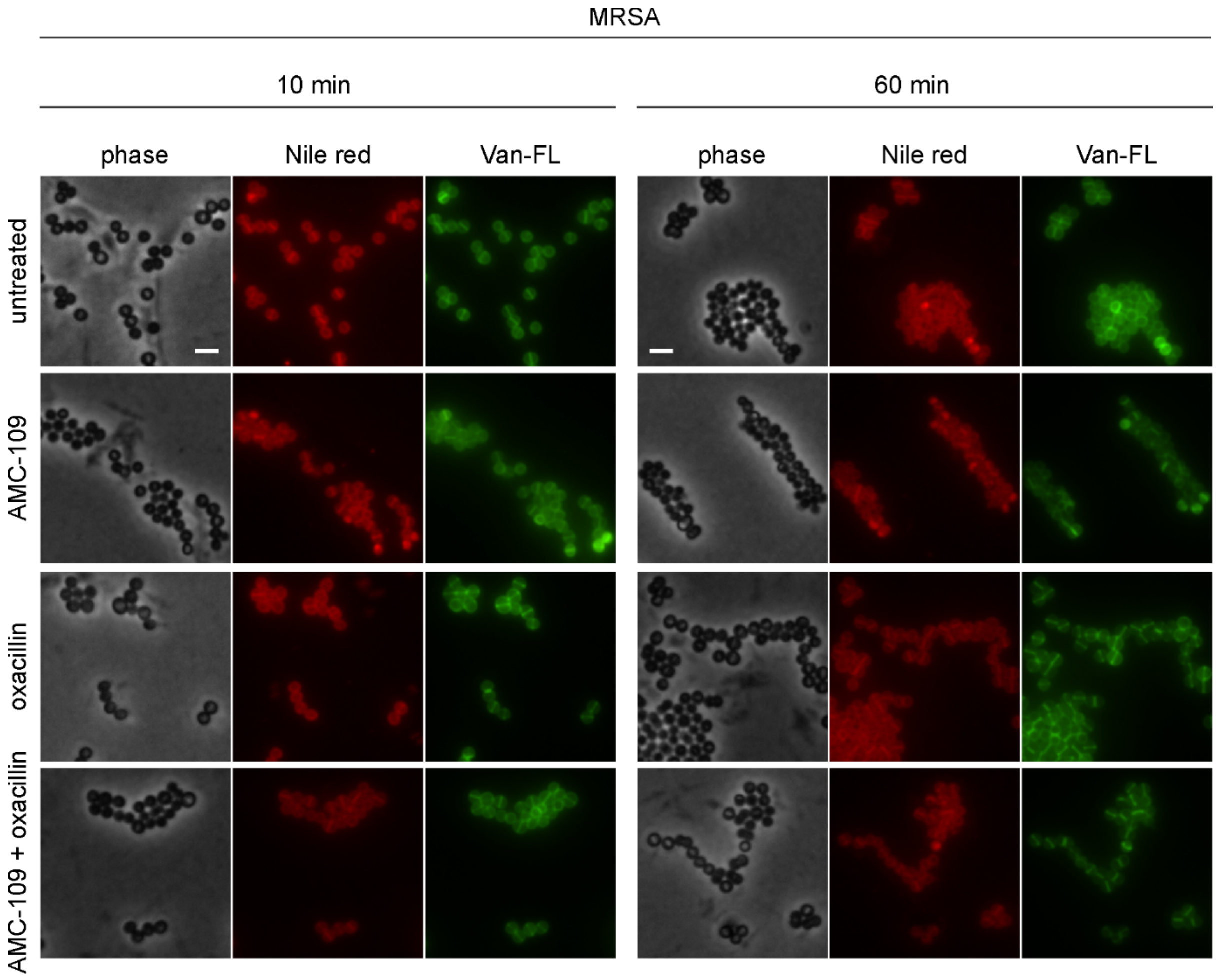


**Figure S9:** Microscopy images of phase contrast, Nile red, and Van-FL staining of MRSA. Nile red is a membrane dye that unselectively stains bacterial cell membranes but partitions into the fluid phase upon phase separation. Accumulation of Nile red in bright spots indicates fluid membrane domains. Van-FL is a fluorescently labeled vancomycin derivative that binds to lipid II, indicating its localization and serving as proxy for sites of active cell wall synthesis. MRSA was treated with 2 µg/ml AMC-109, 3 µg/ml oxacillin, 1 µg/ml AMC-109 + 0.25 µg/ml oxacillin, or left untreated. Scale bars represent 2 µm.

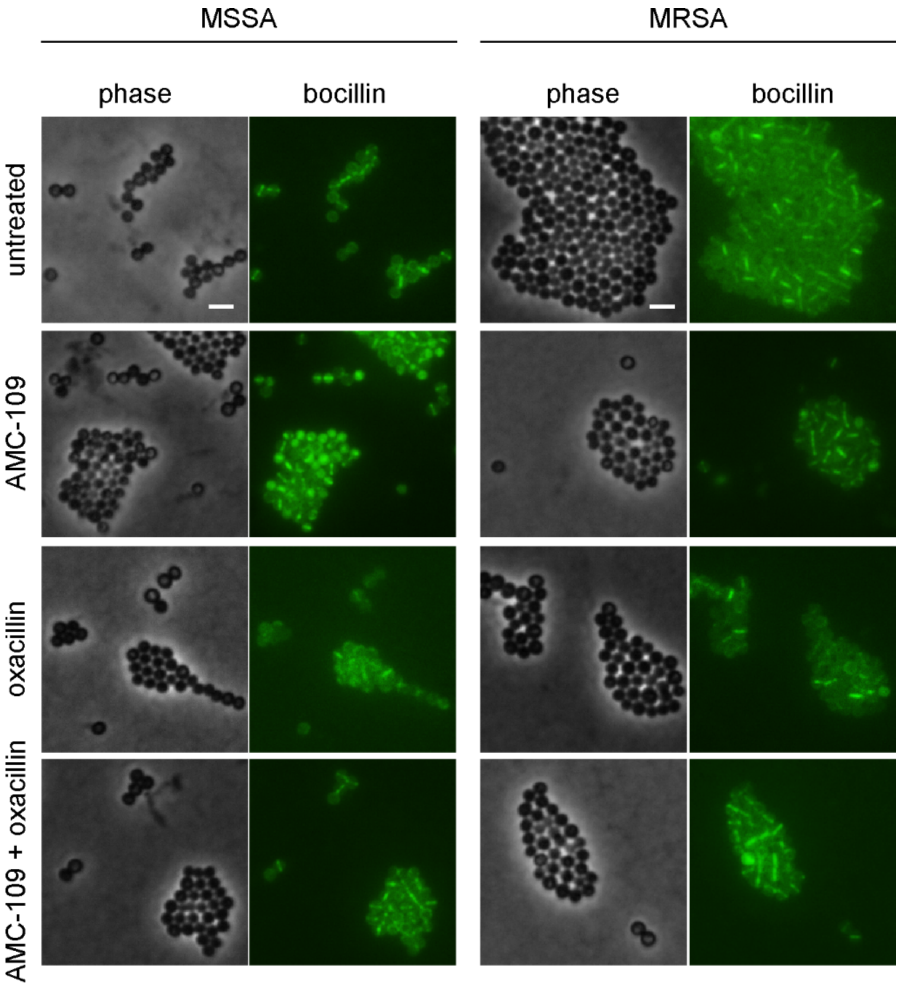


**Figure S10:** Bocillin staining of MSSA and MRSA. Bocillin is a fluorescently labeled penicillin derivative that binds to PBPs visualizing their localization. Images were taken after 10 min of antibiotic treatment. MSSA was treated with 4 µg/ml AMC-109, 0.5 µg/ml oxacillin, 1 µg/ml AMC-109 + 0.0156 µg/ml oxacillin, or left untreated as control. MRSA was treated with 2 µg/ml AMC-109, 3 µg/ml oxacillin, 1 µg/ml AMC-109 + 0.25 µg/ml oxacillin, or left untreated. Scale bars represent 2 µm.

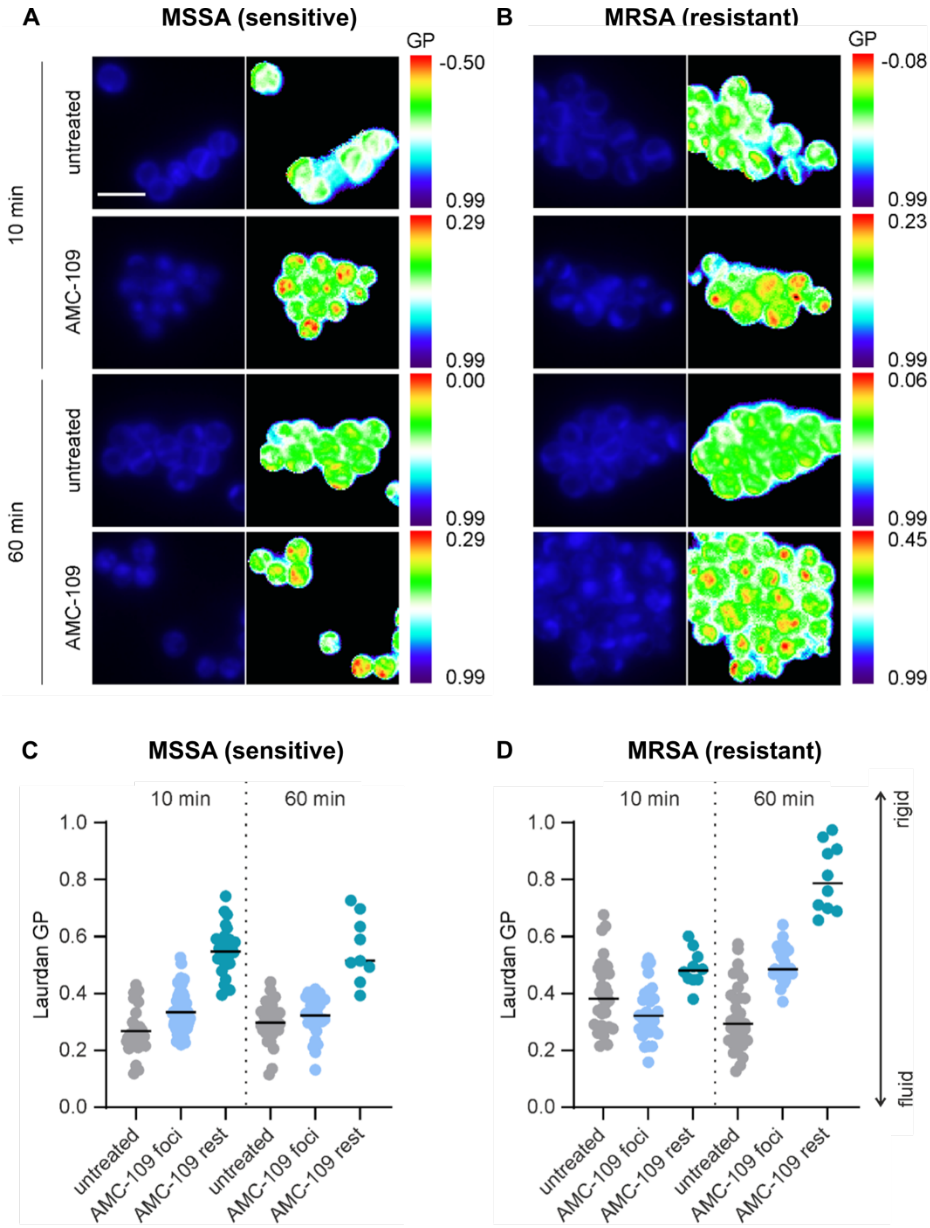


**Figure S11**: Laurdan microscopy images and quantification of (A, C) MSSA and (B, D) MRSA after 10 and 60 minutes of antibiotic exposure. (A-B) Microscopy images showing Laurdan fluorescence at 350 nm excitation and 500 nm emission in blue and Laurdan GP maps calculated based on emission at 460 and 500 nm in rainbow LUT. A higher GP points to more rigid membranes. Scale bar depicts 2 µm. (C-D) Quantification of Laurdan GP measured from individual domains (foci) and out-of-domain (rest) regions.

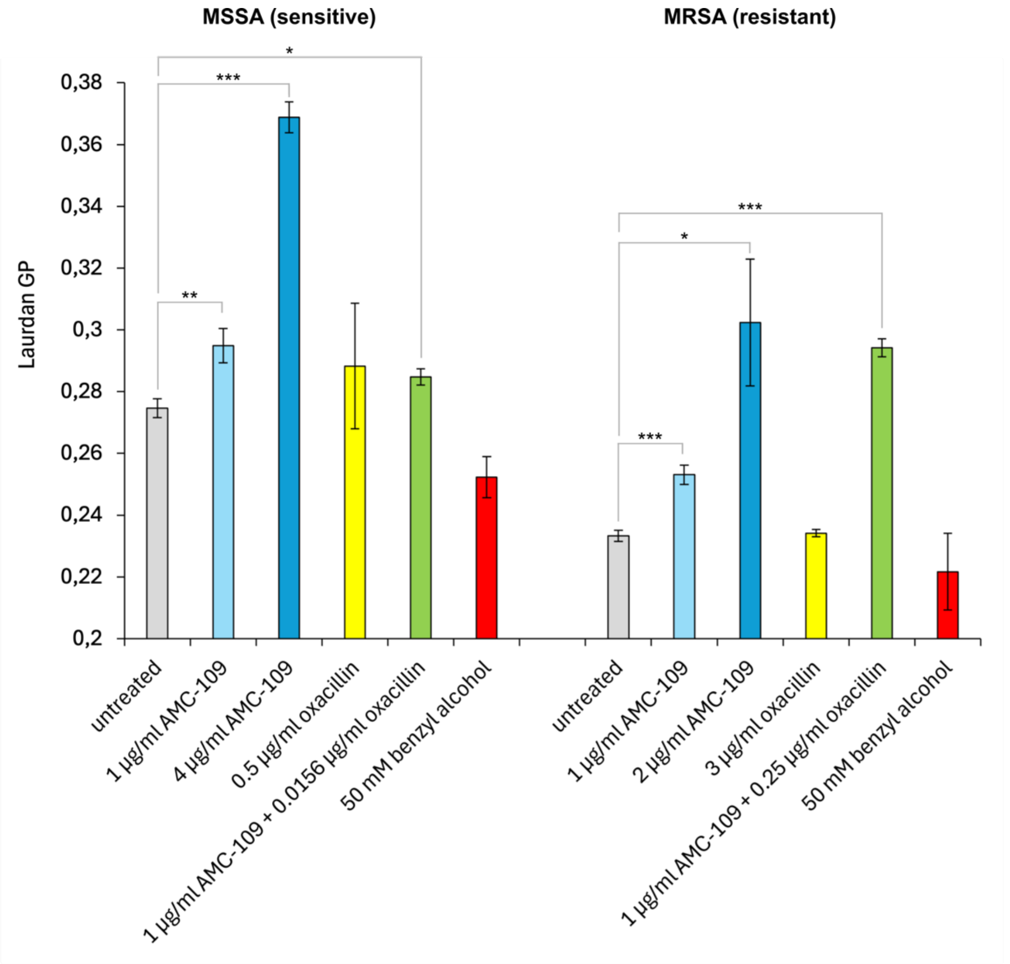


**Figure S12:** Laurdan spectroscopy of batch samples on MRSA and MSSA. Measurements were taken after 10 min of antibiotic treatment or left untreated. Error bars show standard deviation of 3 replicated experiments. Asterisks denote the statistical significance of the difference of the mean Laurdan GP compared to the untreated culture; * indicates p-value $p<0.05$, ** is used for $p<0.01$, and *** for $p<0.001$.

**Table S1**: Comparison of MIC at 24h, 48h, and MBC values in MRSA and MSSA exposed to penicillin G, oxacillin, methicillin, and AMC-109. MIC at 48h coincides with MBC values with the exception of methicillin, which could be explained by poor stability of methicillin over the prolonged time. Maximum values obtained from 5–14 times repeated experiments are shown.

| Bacteria | MRSA | | | MSSA | | |
|---|---|---|---|---|---|---|
| | MIC 24h [µg/ml] | MIC 48h [µg/ml] | MBC [µg/ml] | MIC 24h [µg/ml] | MIC 48h [µg/ml] | MBC [µg/ml] |
| Penicillin G | 2 | 2 | 2 | 0.25 | 0.5 | 0.5 |
| Oxacillin | 2 | 2 | 2 | 0.5 | 0.5 | 0.5 |
| Methicillin | 8 | 32 | 8 | 2 | 4 | 2 |
| AMC-109 | 2 | 2 | 2 | 4 | 4 | 4 |
